# Epigenetic mediation of the onset of reproduction in a songbird

**DOI:** 10.1101/2020.02.01.929968

**Authors:** Melanie Lindner, Veronika N. Laine, Irene Verhagen, Heidi M. Viitaniemi, Marcel E. Visser, Kees van Oers, Arild Husby

## Abstract

Climate change significantly impacts natural populations, particularly phenology traits, like the seasonal onset of reproduction in birds. This impact is mainly via plastic responses in phenology traits to changes in the environment, but the molecular mechanism mediating this plasticity remains elusive. Epigenetic modifications can mediate plasticity and consequently constitute promising candidates for mediating phenology traits. Here, we used genome-wide DNA methylation profiles of individual great tit (*Parus major*) females that we blood sampled repeatedly throughout the breeding season. We demonstrate rapid and directional variation in DNA methylation within the regulatory region of genes known to play key roles in avian reproduction that are in line with observed changes in gene expression in chickens. Our findings provide an important step towards unraveling the molecular mechanism mediating a key life history trait, an essential knowledge-gap for understanding how natural populations may cope with future climate change.

**IMPACT SUMMARY:** Natural populations are increasingly challenged by changing environmental conditions like global increases in temperature. A key way for species to adapt to global warming is via phenotypic plasticity, i.e. the ability to adjust the expression of traits to the environment. We, however, know little about how the environment can interact with an organism’s genetic make-up to shape its trait value. Epigenetic marks are known to vary with the environment and can modulate the expression of traits without any change in the genetic make-up and therefore have the potential to mediate phenotypic plasticity.

To study the role of epigenetics for phenotypic plasticity, we here focus on the great tit (*Parus major*), a species that is strongly affected by global warming and plastic for temperature in an essential phenology trait, the seasonal onset of egg laying. As a first step, we investigated whether great tit females show within-individual and short-term variation in DNA methylation that corresponds to changes in the reproductive state of females. We therefore housed breeding pairs in climate-controlled aviaries to blood sample each female repeatedly throughout the breeding season and used these repeated samples for methylation profiling.

We found rapid and directional variation in DNA methylation at the time females prepared to initiate egg laying that is located within the regulatory region of genes that have previously described functions for avian reproduction. Although future work is needed to establish a causal link between the observed temporal variation in DNA methylation and the onset of reproduction in female great tits, our work highlights the potential role for epigenetic modifications in mediating an essential phenology trait that is sensitive to temperatures.

## INTRODUCTION

Increasing global temperatures have led to shifts in phenology traits of many species over the last decades with major ecological impacts^1,2^. Such shifts in phenology traits include, among many others, leaves unfolding of trees^3^, spring-flowering time of plants^4^, appearance of butterflies^5^, egg laying in seasonally reproducing birds^6^ or the hibernation phenology in squirrels^7^. Species across trophic levels, however, may differ in their phenological sensitivity to climate change and consequently differ in their magnitude of shifts in phenology traits^8^, resulting in phenological mismatches between species of interacting trophic levels that can impact whole ecosystem functioning^9,10^.

A well-documented example is the phenological mismatch between the breeding phenology of great tits (*Parus major*) and the timing of the winter moth (*Operophtera brumata*) caterpillar biomass peak, a key food source of great tit young and hence important determinant of reproductive success^11^. The spring phenology in both species, i.e. laying date in great tits and timing of caterpillar biomass peak, are phenotypically plastic for temperature, but the species differ in their sensitivity to temperature, leading to between-year variation in the magnitude of the phenological mismatch and in the strength of selection on laying date^12^. Individual-based observations from long-term studies allow us to estimate the strength of selection on and the additive genetic variation in laying date components to predict whether laying dates can respond to natural selection^12,13^. Those estimates, however, can vary across environments^14,15^ which complicates such predictions especially if the environment shifts beyond those epreviously observed. Consequently, we might benefit from a more mechanistic understanding of how variation in the environment shapes laying dates which requires insights into the genetic basis of the trait and the molecular mechanism that facilitates the interaction of this genetic basis with the environment. For the seasonal onset of avian reproduction, however, such insights remain elusive^16,17^.

Promising components of the molecular mechanism that mediates laying dates in response to the environment are epigenetic modifications, i.e. chemical modifications of the DNA sequence or chromatin proteins that affect gene expression and consequently trait values without change in the DNA sequence^18^. Interestingly, recent studies in plants^19–21^, insects^22^, and mammals^23,24^ have emphasized the potential for temporal variation in epigenetic modifications to be involved in mediating the temporal expression of phenology traits across taxa. For example, flowering time in *Arabidopsis* is characterized by variation in histone methylation of flowering locus C (*FLC*)^19^, photoperiodic diapause in a parasitic wasp (*Nasonia vitripennis*) is associated with variation in DNA methylation induced by different photoperiods^22^, and gonadal regression in Siberian hamsters (*Phodopus sungorus*) is accompanied by photoperiodically-induced and reversible variation in DNA methylation of type III deiodinase (*dio3*), a gene involved in the photoperiodic regulation of reproduction^23^.

Here, we examine whether short-term variation in DNA methylation mediates the onset of reproduction in a wild songbird species, the great tit. We housed breeding pairs in climate-controlled aviaries, repeatedly blood sampled females throughout the breeding season, and used isolated red blood cells for reduced representation bisulfite sequencing (RRBS) to assess within-individual patterns of temporal variation in DNA methylation. Using two different analytical approaches, we demonstrate rapid and directional variation in DNA methylation at CpG sites within the regulatory region of two genes previously reported in the context of avian reproduction. Our study, therefore, indicates a role of DNA methylation in the molecular mechanism that mediates the seasonal onset of reproduction in a wild songbird species.

## RESULTS

### Differential methylation analysis

We divided the samples into four reproductive stages, based on the reproductive state, i.e. the sampling date centered by the laying date, (Fig. 1) and tested for variation in CpG site methylation in any of the pairwise contrasts between the four reproductive stages.

**Fig. 1.**
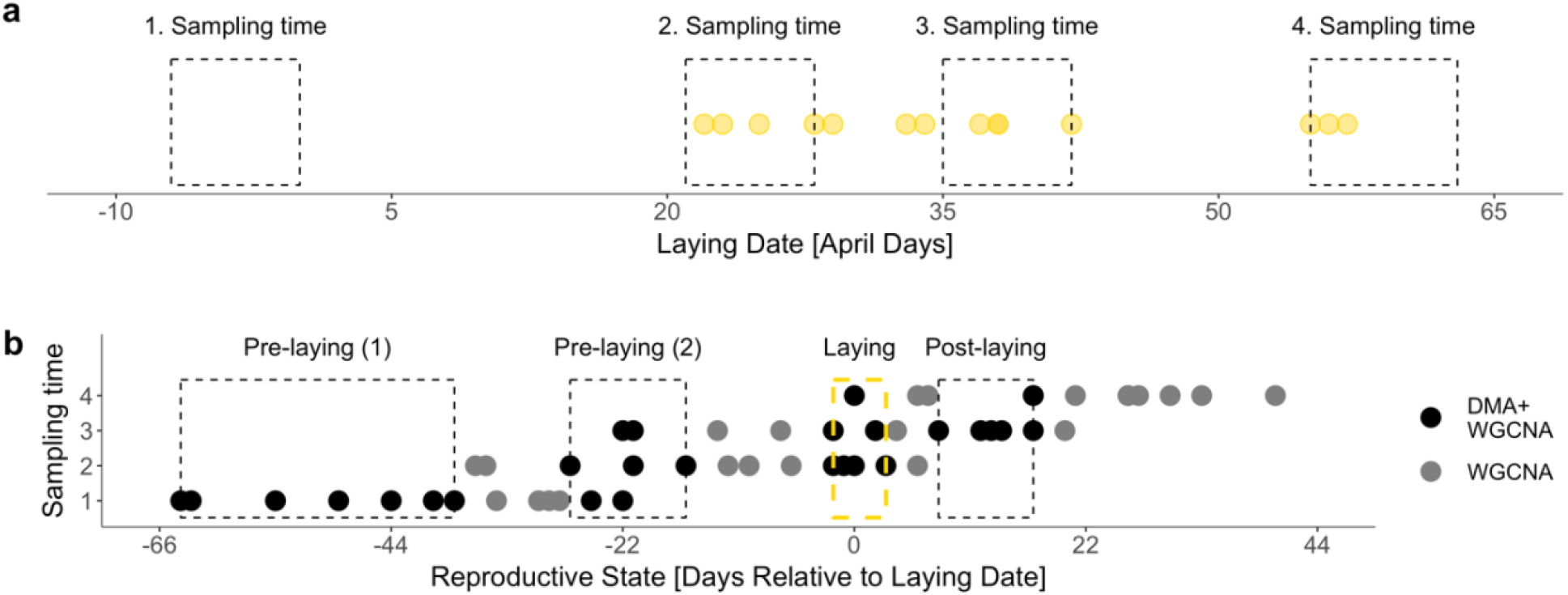
Sampling design. (**a**) Distribution of laying dates with each symbol corresponding to the laying date of one female (n=14). Symbols are transparent (α = 0.5) such that symbols displaying females with overlapping laying dates appear darker. (**b**) The reproductive state (x-axis) separated by sampling time (y-axis) for each sample (n=55). The reproductive state is calculated as the sampling time centered by the respective female’s laying date. Black symbols represent samples used in both analyses, i.e. differential methylation (DMA) and co-methylation analysis (WGCNA), and grey symbols represent samples used in co-methylation analysis only. Groups of samples highlighted in black (separated by samples displayed in grey) represent the four reproductive stages: the first pre-laying, second pre-laying, laying, and post-laying stage (from left to right). Yellow rectangle highlights the reproductive state that corresponds to the laying stage, i.e. samples that were taken at or very close to the respective female’s laying date.

In a genome-wide differential methylation analysis on 5,097 CpG sites within the regulatory region of genes, a region known to affect gene expression in great tits^25^, we found significant variation in CpG site methylation for the different pairwise contrasts (fig. S1), with most of the variation taking place between the pre-laying and post-laying stages. We therefore focus on the comparison between the second pre-laying and post-laying stage (Fig. 2, table S1).

**Fig. 2.**
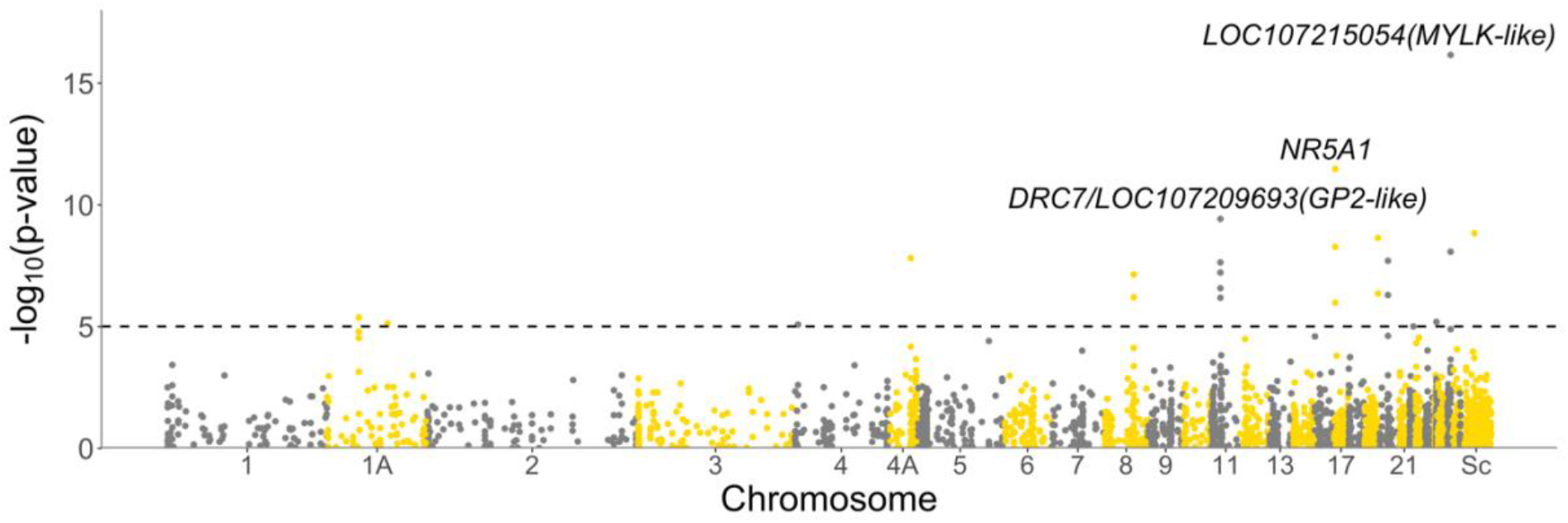
Manhattan plot of p-values (in −log_10_ scale) corresponding to the significance of the variation in DNA methylation between the second pre-laying and post-laying stage. Each tested CpG site (n=5,097) is displayed as dot and plotted against the location of the associated CpG site within the genome. Dotted line marks the genome wide significance threshold (Bonferroni corrected, α_BF_ = −log_10_(0.05/5,097) = 5.01). Alternating colors help to differentiate adjacently displayed chromosomes. ‘Sc’ refers to unplaced scaffolds.

CpG sites that showed the most significant variation in DNA methylation were located within the regulatory region of *LOC107215054* predicted as myosin light chain kinase, smooth muscle-like (*MYLK*-like, p < 6.97e-^17^, Δ Methylation Level = −46 %), the steroidogenic factor 1 (*NR5A1*, p < 3.32e-^12^, Δ Methylation Level = 49 %), the dynein regulatory complex subunit 7 (*DRC7*) and *LOC107209693* predicted as pancreatic secretory granule membrane major glycoprotein GP2-like (*GP2*-like, both p < 3.77e-^10^, Δ Methylation Level = 42 %, table S2).

### Co-methylation analysis

While the differential methylation analysis required *a priori* defined comparisons between groups, we additionally used an unsupervised approach using a weighted co-methylation network analysis (co-methylation analysis), to cluster CpG sites based on the similarity of their methylation profiles across all samples. We found that 2,347 out of 5,097 CpG sites clustered into nine different modules (table S3). We then tested for a correlation between each module’s eigensite, i.e. first principle component, and the reproductive state and found a significant correlation for two modules’ eigensite (Fig. 3).

**Fig. 3.**
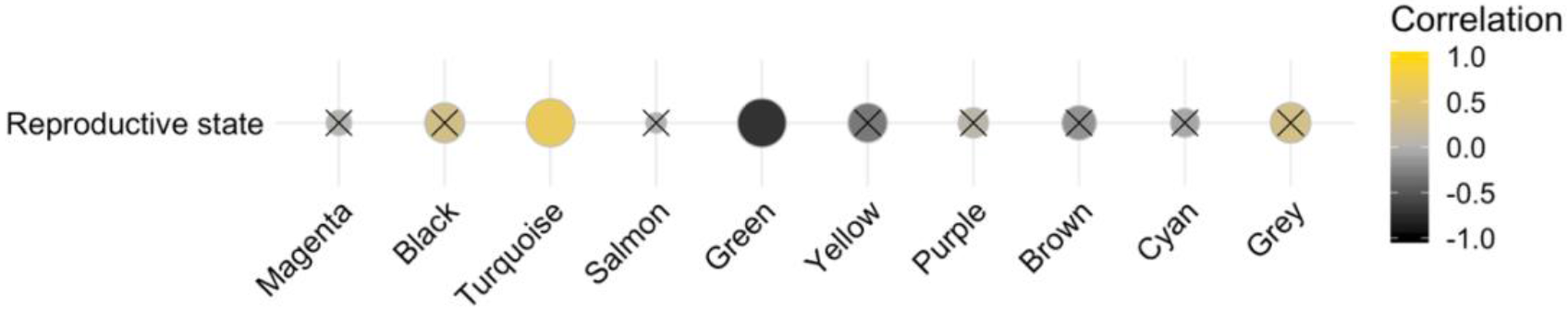
Correlation of the reproductive state with the module eigensite of each detected module. Yellow spheres show a positive correlation, grey spheres no correlation, and black spheres a negative correlation. Size of spheres relates to significance of correlation (the bigger sphere, the lower p-value). Non-significant correlations (p-vale > 0.^001^, Bonferroni-corrected α-threshold) are crossed.

The correlation was positive for the turquoise module (0.66, p = 1.57e-06) and negative for the green module (−0.71, p = 4.82e-08, fig. S2). The turquoise and green module included 18 CpG sites within seven genes and 28 CpG sites within nine genes with significant module membership (MM) and trait-based site significance (SS) (tables S4-S5). The CpG sites with the highest trait-based site significance were located within the regulatory region of *NR5A1* (MM = 0.70, MM p = 3.08e-09, SS = 0.78, SS p < 1.40e-12, turquoise module), and *LOC107213450* (a protein coding but uncharacterized gene, MM = 0.82, MM p = 2.58e-14, SS = −0.81, SS p = 1.28e-13, green module) and *LOC107215054* (*MYLK*-like) (MM = 0.89, MM p = 3.68e-20, SS = −0.75, SS p < 3.41e-11, green module, tables S6-S7).

### Functional analysis

We performed a gene ontology (GO) and STRING analyses with genes found in the differential and co-methylation analyses, but did not detect any significantly enriched GO terms and protein-protein interaction networks. Hence, genes identified here have not been previously described to share biological functions or interact with each other.

### Methylation profiles of most significant findings

To identify the genes that most significantly co-varied with the reproductive state in their DNA methylation profile, we combined the findings from the differential methylation and the co-methylation analysis and found eight genes with one or more CpG sites that showed significant variation in DNA methylation in both analyses (intersections in fig. S3, table S8). We found four of those genes with the differential methylation analysis and the turquoise module of the co-methylation analysis, and five genes with the differential methylation analysis and the green module of the co-methylation analysis. The most significant genes were *NR5A1* and *LOC107215054* (*MYLK*-like). The three CpG sites within the regulatory region of *NR5A1* showed hyper-methylation and the two CpG sites within the regulatory region of *LOC107215054* (*MYLK*-like) showed hypo-methylation at the time females initiated egg laying (Fig. 4), indicative for an epigenetic down-and upregulation of the respective gene.

**Fig. 4.**
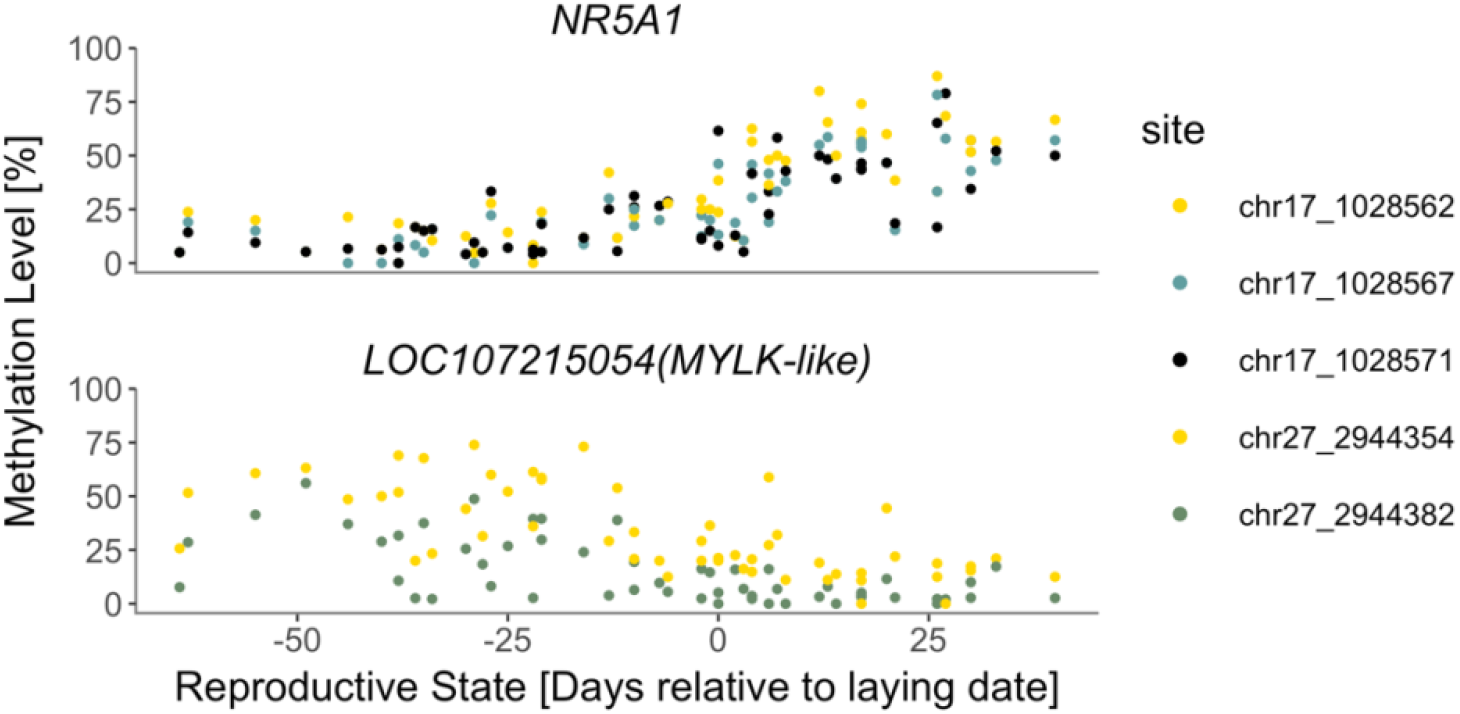
Methylation profile across samples (n=55) of CpG sites within the regulatory region of *NR5A1* (chr17) and *LOC107215054* (*MYLK*-like, chr27) in relation to the reproductive state, i.e. sampling date centered by the respective female’s laying date.

## DISCUSSION

The molecular mechanism that mediates the temporal expression of phenology traits in response to the environment remains largely unknown^16^, but recent studies on plants^19–21^, insects^22^, and mammals^23,24^ emphasize the potential for epigenetic modifications, such as DNA methylation, to be involved in this mechanism. Here, we expand on this by indicating a role of DNA methylation in mediating the onset of reproduction in a wild songbird species, the great tit.

We found two genes, *NR5A1* and *LOC107215054* (*MYLK*-like) that most significantly co-varied with the reproductive state in their DNA methylation profile, when we combined findings from the differential methylation and the co-methylation analysis. *NR5A1* encodes for a transcription factor that regulates the expression of many important genes within all levels of the reproductive axis^26,27^ and most genes involved in gonadal steroidogenesis^28^. For example, *NR5A1* modulates the expression of the steroidogenic acute response protein (*STAR)* that transfers cholesterol to initiate an enzymatic cascade that comprises steroid synthesis, essential for folliculogenesis^29^. *NR5A1* is expressed in the major steroidogenic tissues, such as theca and granulosa cells in the ovary^27,30,31^ or hypothalamus (e.g. regulating hypothalamic pituitary gonadotroph organization and function^32^). Consequently, *NR5A1* plays a key role in female reproduction such as ovarian functioning^26,33^ and steroidogenesis^34^. Expression studies of *NR5A1* in chickens are concordant with the change in DNA methylation observed here as *NR5A1* is up-regulated during egg laying relative to brooding^35^. This expression pattern, in combination with the essential role of *NR5A1* in ovarian functioning and steroidogenesis, suggests that *NR5A1* acts as a key transcription factor in the initiation and upkeep of the reproductive cascade that mediates egg laying.

*LOC107215054* (*MYLK*-like) is predicted as myosin light chain kinase, smooth muscle-like and shares sequence regions with the myosin light chain kinase (*MYLK*), a kinase that facilitates phosphorylation of the myosin light chain, a process essential for shell gland contractile activity during oviposition, i.e. egg laying^36,37^. In chicken ovaries *MYLK* was up-regulated during egg laying^38^, again a pattern consistent with the decrease in DNA methylation we observed at *LOC107215054* (*MYLK*-like). Whether both genes function as myosin light chain kinases remains, however, to be established. The methylation profiles of other genes with CpG sites that show significant variation in DNA methylation in both analyses are similar to those described above (fig. S4) and their functions are described in the supplementary discussion S1.

Our results indicate a functional role for DNA methylation in the molecular mechanism that mediates the seasonal onset of reproduction in great tits. This indicated link is based on CpG sites that significantly vary in DNA methylation with the reproductive state and well-described functions for avian reproduction of the genes in which those CpG sites are located. Also, the methylation patterns found here are supported by multiple CpG sites within the regulatory region of genes and the methylation patterns match with expression patterns of the respective genes in chickens^35,38^. Nevertheless, future studies would be needed to experimentally establish a functional link between observed methylation patterns and the seasonal onset of reproduction.

We used red blood cells from repeatedly sampled great tit females to examine temporal and genome-wide DNA methylation patterns. The strength of this approach is that we are able to examine how DNA methylation within an individual varies with the reproductive state. More informative tissues such as gonads, hypothalamus, or liver cannot be sampled repeatedly^39^ and do not allow a direct correlation with the reproductive state as some females would be sampled before egg laying. The biological relevance of red blood cells as informative tissue with respect to the seasonal onset of reproduction, however, remains unclear. Recent evidence from great tits suggests that red blood cell and liver methylation changes predictably in both a tissue-specific and a tissue-general manner^40,41^. Hence, observed patterns in red blood cell methylation may be established in a tissue-general as well as tissue-specific manner and thus be informative about DNA methylation patterns in other tissues, although the generality of these findings needs to be established.

We here demonstrate short-term variation in CpG site methylation within the regulatory region of genes with known function for avian reproduction. Although future work is needed to establish a causal link between the observed temporal variation in DNA methylation and the seasonal onset of reproduction, our work highlights the potential role for an epigenetic modification to play an important role in the molecular mechanism that mediates the environmental sensitivity of phenology traits.

## METHODS

### Study system

We used the great tit, a well-known model species in ecology and evolution with a reference genome^25^, SNP arrays (10k^42^ and 650k^43^), and whole transcriptome and methylome for various tissues^25,41,44^. The individuals included are part of a bidirectional selection experiment for early and late reproduction using genomic selection^45,46^ (details in methods S1). For the experiment, we housed 36 breeding pairs of the F2 generation in climate-controlled aviaries mimicking natural temperature and photoperiod patterns of a cold and warm year in the Netherlands^47,48^.

### Blood sampling and sample selection

Pairs were blood sampled biweekly from January to July between 08:30 AM and 14:30 PM from the jugular vein (up to 150 μl) and for DNA extraction red blood cells were separated from the plasma^49^ (details in methods S2). We chose samples from 16 females of the early selection line collected at four sampling times based on the females’ realized laying dates (Fig. 1): the day when (1) day length > 12h, (2) 25% of the females exposed to the warm temperature environment had initiated egg laying, (3) 25% and 50% of the females exposed to the cold and warm temperature environment respectively had initiated egg laying, and (4) 50% of the females exposed to the cold temperature environment had initiated egg laying). One blood sample is missing of one female (due to the female incubating) at the fourth sampling time, resulting in a total of 63 samples.

### Sample processing and reduced representation bisulfite sequencing

Library preparation and sequencing was done by the Roy J. Carver Biotechnology Centre (University of Illinois at Urbana-Champaign, USA). For details on sample processing and sequencing see respective publication^49^. RRBS data have been submitted to the NCBI BioProject database (http://www.ncbi.nlm.nih.gov/bioproject/) under BioProject PRJNA208335 and accession number SRX3209916-SRX3209919.

### Sequence alignment and CpG site calling

Sequence alignment^49^, CpG site calling^49^, quantification of DNA methylation^49^ and general methylation statistics of the data set^50^ are described in respective publications. CpG sites with a minimum of 10x coverage in all samples were included for statistical analysis resulting in methylation information of 522,645 CpG sites in each of the 63 samples. We excluded samples collected from two females (one from the warm and one from the cold temperature environment) as these females did not initiate egg laying during the experiment, reducing the data set by eight samples.

### Regulatory region and gene annotation

We defined a region spanning 2000 bp upstream to 200 bp downstream of the transcription starting site as the regulatory region of a gene to annotate identified CpG sites using the *Parus major* reference genome build 1.1 (https://www.ncbi.nlm.nih.gov/assembly/GCF_001522545.2) and R packages GenomicFeatures^51^ v1.30.0 and rtracklayer^52^ v1.42.2. We found 223,282 CpG sites located within the regulatory region of 12,325 genes (out of 18,611 annotated genes) using BEDtools^53^ v2.26.0. Please note that one site can be associated to the regulatory region of two genes if the two genes are located on opposite strands and have overlapping regulatory regions.

### Differential methylation analysis

We calculated the reproductive state as the sampling date centered by the respective females’ laying date and used this to group samples into four different reproductive stages: first pre-laying stage (64 to 38 days prior to laying), second pre-laying stage (27 to 16 days prior to laying), laying stage (2 before to 3 days after laying), and post-laying stage (8 to 17 days after laying, Fig. 1). Each stage constitutes of seven samples (making a total of 28 samples) and samples of all 14 females are present in at least one stage. We filtered CpG sites such that only sites with at least 10% change in average methylation level between any of the four reproductive stages^54^ are included in the differential methylation analysis, reducing the data set to 5097 CpG sites.

We applied a Generalized linear mixed model (GLMM) approach to identify sites that were significantly differentially methylated between any of the four reproductive stages using R package lme4qtl^55^ v0.1.10. Thus, for each of the 5097 CpG sites we fitted a GLMM with binomially distributed errors as specified in equation (1)

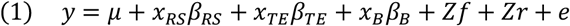

with *y* for the modeled response, a two-column matrix of methylated and unmethylated counts as dependent variable to account for variation in CpG site coverage^56,57^, *μ* for the intercept term, *x_RS_*, *x_TE_*, and *x_B_* for the vector relating individuals to their reproductive stage, temperature environment, and batch, respectively, *β_RS_*, *β_TE_*, and *β_B_* for the reproductive stage, temperature environment, and batch effect, respectively, *Z* for an incidence matrix relating individuals to their observed values, *f* and *r* for the random effects for repeated measurements and relatedness, and *e* for the residuals. Genomic relatedness was calculated using R package GenABEL^58^ v1.8.0. Control parameters used were boundary distance of 0.01, maximum number of iterations of 2×10^8^, and “bobyqa” as an optimizer to speed up computation and optimize convergence^59^. For each CpG site we used the fitted model to infer the estimated marginal means (EMMs) for each level of the reproductive stage and tested for an effect between any pair-wise combinations of EMMs using R package emmeans^60^ v1.3.3. For all pair-wise comparisons we observed an underlying uniform distribution that increased towards low p-values and peaks at p-value > 0.1 (fig. S5). For each GLMM (i.e. each CpG site) we calculated the dispersion statistic following Zuur et al. (2013) and removed highly under or over-dispersed models (keeping only models with dispersion statistic within the range of 0.7 to 1.3). Dispersion statistics were normally distributed with a median of 1.09 (95% CI [1.08, 1.10], fig. S6). We used a Bonferroni corrected α-threshold (p-value < 9.81e-06) to accept the variation in DNA methylation of a CpG site between the respective reproductive stages as significant.

### Co-methylation analysis

For the weighted co-methylation network analysis (co-methylation analysis) we used R package WGCNA62 v1.66 to cluster CpG sites into modules based on similarity in methylation pattern throughout the reproductive state^63–67^. In contrast to the differential methylation analysis, all samples of females that have initiated egg laying were used. The same set of 5,097 CpG sites as for the differential methylation analysis was used for network construction and model detection with a color assigned to each detected module (details in methods S3). We summarized the methylation profile of each module as the module eigensite equivalent to the first principal component of a module based on the methylation profile of all CpG sites within that module (see fig. S7 for variance explained by the first to tenth principle component of each detected module). For each detected module, we tested whether the methylation profile of CpG sites within a module co-varied with the reproductive state by correlating the module eigensite of a module with the reproductive state and examined the associated p-value (details in methods S4). Next to the reproductive state, we also tested for a correlation of module eigensites with other sampling traits (i.e. laying date *per se*, sampling date, female id, and temperature environment, results are presented in fig. S8 and table S9). We accepted a correlation with p-value < 0.001 (Bonferroni-corrected α-threshold, based on number of modules and traits tested) as a significant correlation. Modules with a significant correlation to the reproductive stage (here turquoise and green modules, Fig. 3 and fig. S2) and CpG sites within such modules that show a significant module membership and a significant trait-based site significance are potential candidates for further validation^62^ (see methods S5 for how these were estimated). We accepted the module membership and trait-specific site significance as significant if p < 6.05e-05 and p < 2.14e-04 (Bonferroni-corrected α-threshold, based on number of sites within each module) for a CpG sites with a the turquoise and green module, respectively. We ranked CpG sites within each module based on their p-value for the trait-based site significance.

### Functional analysis

We performed gene ontology (GO) analyses for genes identifies with the differential methylation analysis and the co-methylation analysis (for details see Supplementary Methods 6 and Supplementary Table 10) using ClueGO^68^ v2.5.3 plug-in for Cytoscape^69^ v3.7.1. We used the human annotations, GO categories ‘biological process’, ‘cellular components’, ‘molecular functions’ and KEGG pathways (versions from 09.07.2019), and a custom background lists of all genes with a CpG site in their regulatory region. We specified selection criteria for GO terms such that >5% of the genes associated with a GO term and >3 genes associated with the GO term had to be present in input genes. For the enrichment and functional analyses we used a two-sided enrichment/depletion test, p-value correction for multiple testing via Bonferroni step down, and set the network specificity to ‘medium’ ranging from third to tenth GO level. In addition to the GO analyses, we used the STRING^70^ v1.4.2 plug-in for Cytoscape^69^ v3.7.1 to construct protein-protein interaction network analyses for the same genes as in GO analyses and used a confidence cutoff to 0.7.

### Methylation profiles of most significant findings

For the differential methylation analysis, we ranked genes based on the p-value of the CpG site that showed the most significant variation in DNA methylation between the second pre-laying and post-laying stage (table S2). For the co-methylation analysis, we ranked genes for the turquoise and green module separately such that the rank is based on the p-value for the trait-specific site significance of the CpG site that showed the highest trait-specific site significance within a gene (tables S6-S7). We calculated the sum of ranks (i.e. sum of the rank given based on the differential methylation analysis and the rank given based on the co-methylation analysis, table S8) for all genes with a CpG site that is significant in both analyses. Next to exploring the methylation profiles of CpG sites within the regulatory region of the two top ranked genes, we explored the methylation profile of all remaining CpG sites that were significant in both analyses (results presented in fig. S5), significant in the differential methylation analysis but not in the weighted co-methylation network analysis (results presented in fig. S9), and significant in the weighted co-methylation network analysis but not in the differential methylation analysis (results for the turquoise and green module are presented in fig. S10 and fig. S11, respectively).

## Supporting information

Supplemental R script: Co-Methylation Analysis

Supplemental R script: Differential Methylation Analysis

## DATA AVAILABILITY

RRBS data have been submitted to the NCBI BioProject database (http://www.ncbi.nlm.nih.gov/bioproject/) under BioProject PRJNA208335 and accession numbers SRX3209916-SRX3209919.

## CODE AVAILABILITY

R code for the differential methylation and co-methylation analyses are available as supplementary R scripts.

## NOTES

## Acknowledgments

We thank Koen Verhoeven for commenting on earlier drafts of the manuscript, Christa Mateman and colleagues at NIOO-KNAW for lab assistance, Bart van Lith and Ruben de Wit for assistance during the experiments, Jeroen Laurens and Gilles Wijlhuizen for technical assistance prior and during the experiments, and the animal care takers at the NIOO-KNAW for the care of the birds. This work was supported by the Research Council of Norway Centre through its Centre of Excellence funding (223257), a personal grant to A.H. (239974) from the Norwegian Research Council and a European Research Council (ERC-2013-AdG 339092) to M.E.V.

## Author contributions

A.H., M.E.V., and K.v.O. designed the experiment, I.V. conducted the experiments, H.M.V. conducted the sequences alignment, and M.L. conducted the statistical analysis with the help of V.N.L. M.L. and A.H. wrote the first draft of the manuscript with input from all authors.

## Materials and correspondence

Correspondence and requests for materials should be addressed to (ML) or (AH).

## Ethics approval

This study was performed with all relevant ethical regulations and under the approval by the Animal Experimentation Committee (DEC), Amsterdam, The Netherlands, protocol NIOO 14.10 and addendum 2 to this protocol.

