## Supplemental Material for "Epigenetic mediation of the onset of reproduction in a songbird"

### SUPPLEMENTARY DISCUSSION

*Discussion S1. LOC107209693* is predicted as predicted as pancreatic secretory granule membrane major glycoprotein GP2-like (*GP2*-like*)* which is a component of the inner perivitelline layer surrounding the avian ovum during ovulation that helps to maintain the structural integrity of the vitelline membrane[34,35]. The inner perivitelline layer mechanically supports the ovum and mediates the initial interaction between the ovum and spermatozoa in which glycopeptides are suggested to play an essential role[35]. Consequently, *GP2* might have a function in fertilization, but whether both genes, i.e. *GP2*-like and *GP2*, share their biological function remains to be established. Furthermore, *GP2* was reported as a modulator for immune response[36], e.g. by binding pathogenic enterobacteria[37].

The dynein regulatory complex subunit 7 (*DRC7*) and the cilia and flagella associated protein 45 (*CFAP45*) both regulate cilia and flagella motor activity[38–40]. In eukaryotic organisms, e.g. humans[38] or mice[41], cilia are found on the surface most cell types and play essential roles in all life stages ranging from embryonic development to reproduction. In mammals, for example, the multi-ciliated cells within the infundibulum generate a current that draws the ovulated oocyte into the oviduct[42]. Whether this is the case in female birds remains, to our knowledge, unexplored.

The solute carrier family 6 comprises transporters with essential roles in neurotransmission, cellular and whole body homeostasis[43]. The solute carrier family 6 member 9 (*SLC6A9*) transports the amino acid glycine and is distributed widely in human organs. A specific function for *SLC6A9* in avian reproduction remains, to our knowledge, unexplored.

The disks large homolog 3 (*DLG3*) is a protein concentrated in excitatory synapses in which it mediates synaptic transmission via binding to the NMDA receptor[44]. *DLG3* is suggested to play an important role in synaptic development and plasticity[44,45], but a specific function for *DLG3* in avian reproduction remains, to our knowledge, unexplored.

The clustered mitochondria protein homolog (*CLUH*) is a protein involved in controlling cell energetic and metabolic status via regulation of mitochondrial distribution and biogenesis[46]. A specific function for *CLUH* in avian reproduction remains, to our knowledge, unexplored.

### SUPPLEMENTARY METHODS

*Methods S1.* In the artificial selection experiment great tis were genetically selected for either early or late reproduction via genomic selection. More detailed description of the genomic selection[6] and selection line experiment[7] can be found in respective publications. The selection criterion for genomic selection is the 'genetic breeding value' (GEBVs), an estimate corresponding to an individual's value for the trait under selection based on its combination of 'single nucleotide polymorphisms' (SNPs). We established GEBVs with the 'genomic best linear unbiased prediction' (GBLUP) approach[32] using a 'training population' of 2015 females for which lay dates (i.e. the day on which a female has laid its first egg of the year) and genotypes using the Great Tit Affymetrix 650k SNP chip[3] were available. Once established, GEBVs can be estimated without information on the phenotype (i.e. lay date) which speeds up the selection procedure as (1) breeding pairs can be selected previous to a female's first breeding season and (2) the GEBVs can be estimated for both sexes such that males (that do not express the phenotype) can contribute to the effect of the selection line procedure. For the selection line experiment, we collected the chicks (F1 generation) of wild breeding pairs (parental (P) generation) at day 10 post-hatching in 2014, genotyped the chicks, calculated their GEBVs, and paired them such that birds with the most extremely negative and positive GEBVs were used for the early and late selection line, respectively. The following year breeding pairs of the F1 generation bred in captivity and resulting chicks (F2 generation) underwent the same selection line procedure to breed in the subsequent year (2016). For the experiment we housed 36 breeding pairs of the F2 generation in climate-controlled aviaries mimicking natural temperature and photoperiod patterns of a cold (2013) and warm (2014) year in the Netherlands (realized temperature pattern[8] and daily average and daily minimum and maximum temperatures[9] can be found in respective publications). Aviaries were provided with three nest boxes and nesting material was provided from March onwards. More details on housing conditions[8] can be found in the respective publication.

*Methods S2.* For blood sampling pairs were divided in two batches and each batch was sampled biweekly from January to July. During a sampling day, birds (n=36, 18 pairs randomized by temperature environment and selection line) were sampled between 08:30 AM and 14:30 PM such that every pair was blood sampled from the jugular vein (up to 150 µl) within 10 min after capture. We placed each tube on ice till 12 individuals have been blood sampled to then spin samples in a centrifuge at 14,000 rpm for 12 min for separation of plasma and red blood cells. Afterwards, we removed the plasma using Hamilton syringes (Merck KGaA), resolved the remaining RBCs in Queens buffer, and temporarily stored the tubes at RT before samples were processed. We extracted the DNA from RBC samples using FavorPrepT M 95-well Genomic DNA Kit and quality of the extraction was assessed by running 1% agarose gel electrophoresis and using Nanodrop 2000.

*Methods S3.* For the weighted co-methylation network analysis we used R package WGCNA[23] v1.66. The package implements functions to cluster CpG sites into modules based on similarity in methylation pattern over samples. The modules, in turn, can be correlated to sample traits of interest. The package is mainly applied to expression data[9,23], but is increasingly used for methylation data[24–28]. Here we used all 55 samples of females that initiated egg laying (no outliers detected using hierarchical clustering, fig. S12) and the same set of 5,097 CpG sites as for differential methylation analysis as we found a skewed scale free topology when using all 223,282 CpG sites located within the regulatory region of genes (fig. S13). We used WGCNA::blockwiseModules for network construction and model detection. A network is specified by its adjacency matrix $a_{ij}$ (symmetric $n \times n$ matrix with entries in $[0,1]$) whose component $a_{ij}$ encodes the network connection strength between site $i$ and $j$. The adjacency, see equation (2), is constructed from correlations

$$(2) \quad adjacency = 0.5 * (1 + cor)^{power}$$

with *power* being the soft thresholding power which is used to emphasize strong correlation on the expense of weak correlations. We used WGCNA::pickSoftThreshold to choose the *power* based on the scale free topology criterion[33]. We picked a soft threshold of 14 as this is the lowest power at which scale free topology index $R^2 > 0.9$ (truncated $R^2 = 0.99$) and mean connectivity decreases to 2.50 (fig. S14). Thus, the weighted networks used here are highly robust with regard to the power. Furthermore, weighted networks allow the adjacency to take on continuous values between 0 and 1 (in unweighted networks adjacency is either 1 or 0, which does not reflect the continuous nature of the underlying co-methylation information). We specified a 'mergeCutHeight' of 0.65 in WGCNA::blockwiseModules to prevent high adjacency between module eigensites (fig. S15, see fig. S16 with 'mergeCutHeight' of 0.15).

*Methods S4.* We tested for trait significance of modules by correlating the module eigensites with sample traits (using WGCNA::cor) and generating associated p-values (using WGCNA::corPvalueStudent). Here, we were particularly interested in testing the reproductive state as sample trait, but also tested lay date *per se*, sampling date, female id, and temperature environment. We accepted a correlation with p-value < 0.001 (Bonferroni-corrected $\alpha$-threshold) as significant.

*Methods S5.* CpG sites with high module membership (MM) and trait-based site significance (SS) within such a module are natural candidates for further validation[23]. We used WGCNA::signedKME to calculate the MM (also known as signed eigensite-based connectivity) for each CpG site within modules significantly correlated to the reproductive state. The MM of a CpG site is based on the correlation of the CpG site-specific methylation profile with the module eigensite of the respective module. The trait-based SS of a CpG site is based on the correlation of the CpG site-specific methylation profile with the reproductive state. See fig. S17 for how MM and trait-based SS correlate for each site within the turquoise and green module.

*Methods S6.* We performed GO analyses for genes found in either of both the differentially methylation analysis and the weighted co-methylation network analysis. Gene lists for the differentially methylation analysis included genes with a CpG site that showed significant variation in DNA methylation between the second pre-laying and post-laying stage, and genes with a CpG site that showed significant variation in DNA methylation in any of the pairwise comparisons. Gene lists for the co-methylation analysis included genes with a CpG site that was part of the turquoise module and showed significant MM and trait-based SS, genes with a CpG site that was part of the green module and showed significant MM and trait-based SS, and genes with a CpG site that was part of the either the turquoise or the green module and showed significant MM and trait-based SS (table S10). The costume background list constitutes of all genes with a CpG site in their regulatory region. Please note that some genes within the great tit reference genome[1], were annotated as LOC genes which can be predicted proteins, uncharacterized proteins, or non-coding RNA. As not all LOC genes (especially in the background reference list) could be explored manually for predictions, we excluded all LOC genes from the GO analysis using the custom background list and included LOC genes with known prediction in analysis using the whole human annotation as background.

SUPPLEMENTARY FIGURES

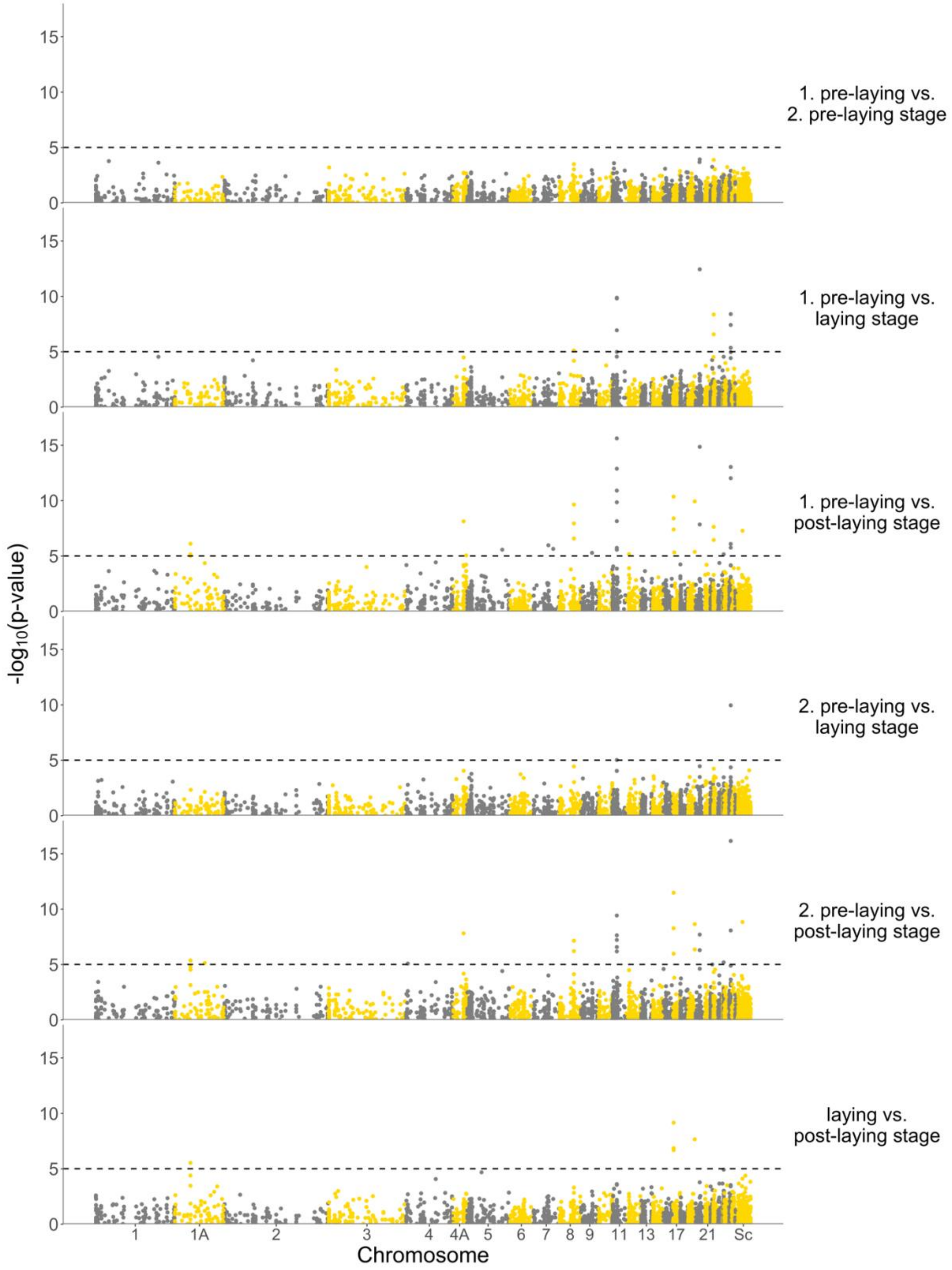


**Fig. S1.** Manhattan plots of CpG sites derived from differential methylation analysis. For each site (displayed as dots) we tested for a difference in methylation level between any of the four reproductive stages (i.e. six pairwise comparisons) using a differential methylation analysis. Each pairwise comparison is displayed in a separate

Manhattan plot (top to bottom) such that every site is included once per pairwise comparison. p-values are displayed as $-\log_{10}$(p-value) and plotted against the location of the associated CpG site within the genome. Dotted line marks the genome wide significance threshold (Bonferroni corrected, $\alpha_{BF} = -log_{10}(0.05/5{,}097) = 5.01$).

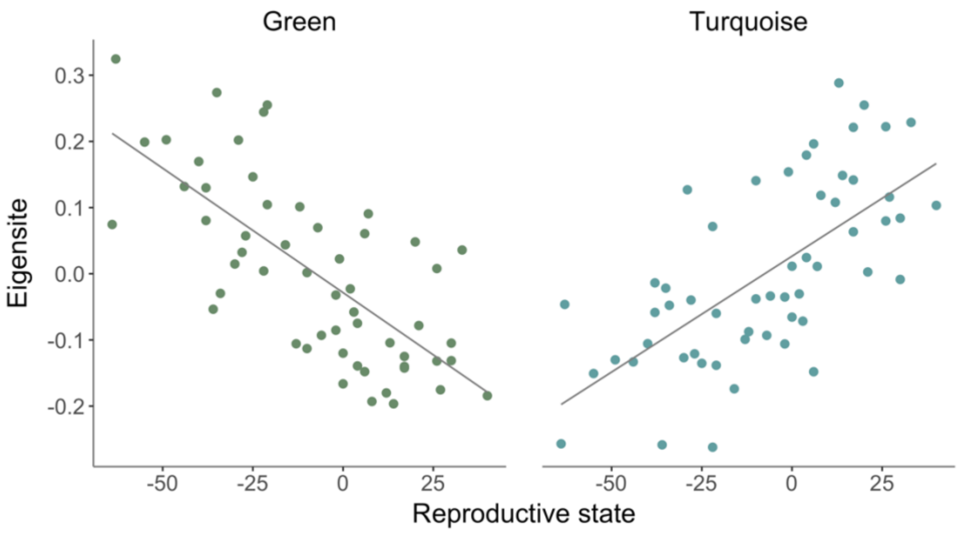


**Fig. S2.** Significant correlations between the module eigensites and the reproductive state. Left panel shows the correlation for the green module (-0.71, p-value = 4.82e-08) and the right panel shows the correlation for the turquoise module (0.66, p-value = 1.57e-06).

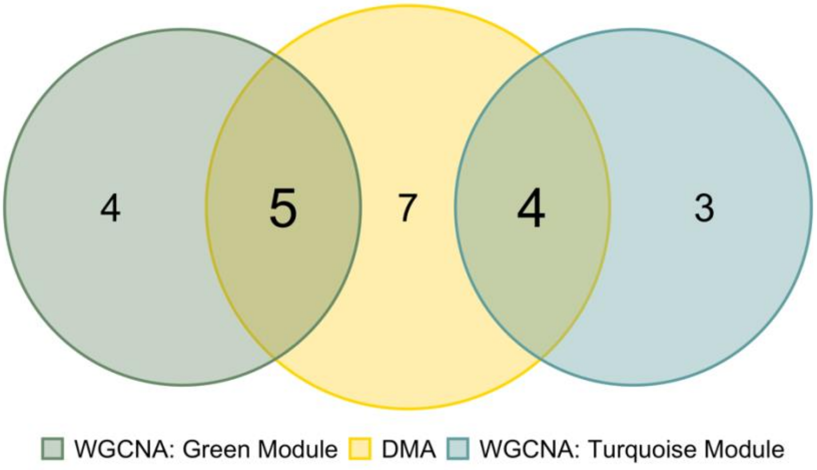


**Fig. S3.** Venn diagram for the combination of the differential methylation analysis and co-methylation analysis. The color of spheres indicates the respective analysis: differential methylation analysis (yellow) and co-methylation analysis (green: green module, blue: turquoise module). Numbers within the intersects relate to genes with CpG sites that show significant variation in DNA methylation in both analyses.

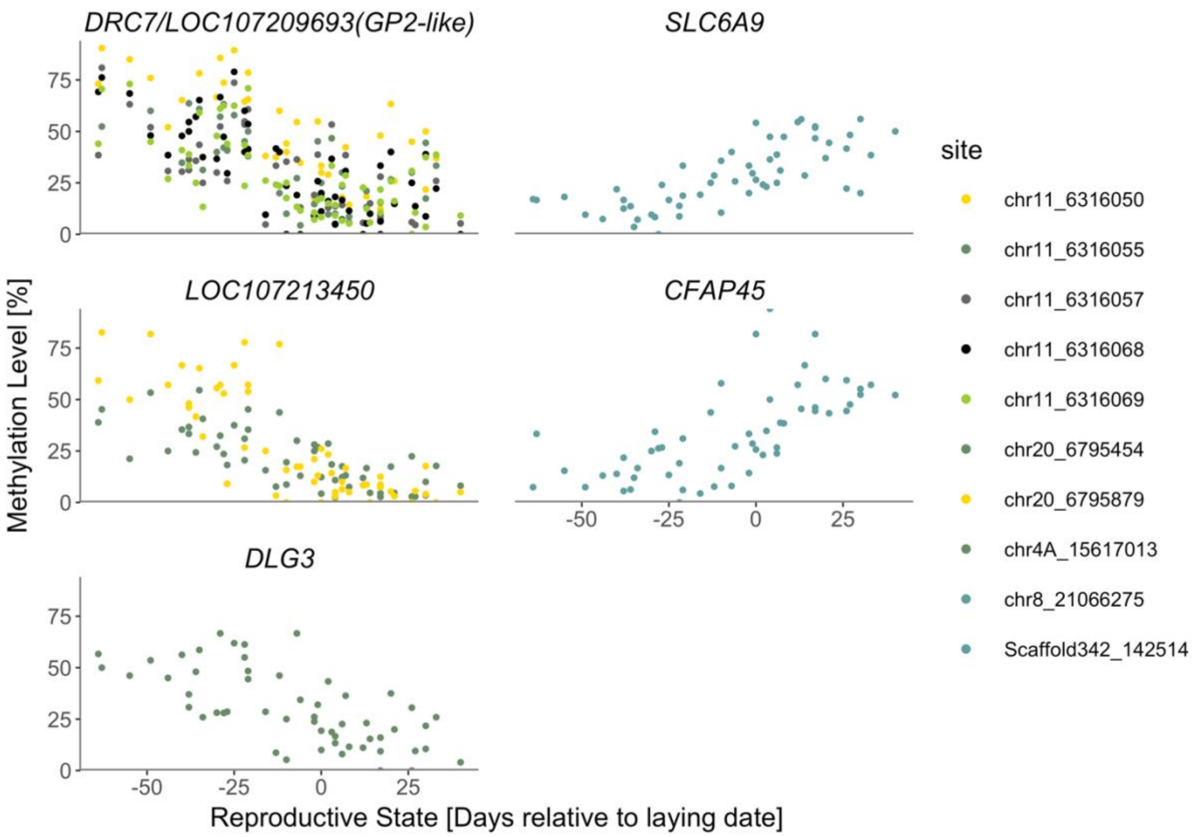


**Fig. S4.** Methylation profile across samples (n=55) of CpG sites within the regulatory region of *DRC7* / *LOC107209693* (*GP2*-like, chr11, green module), *LOC107213450* (chr20, green module), *DLG3* (chr4A, green module), *SLC6A9* (chr8, turquoise module), and *CFAP45* (Scaffold342, turquoise module) in relation to the reproductive state, i.e. sampling date centered by the respective female's laying date.

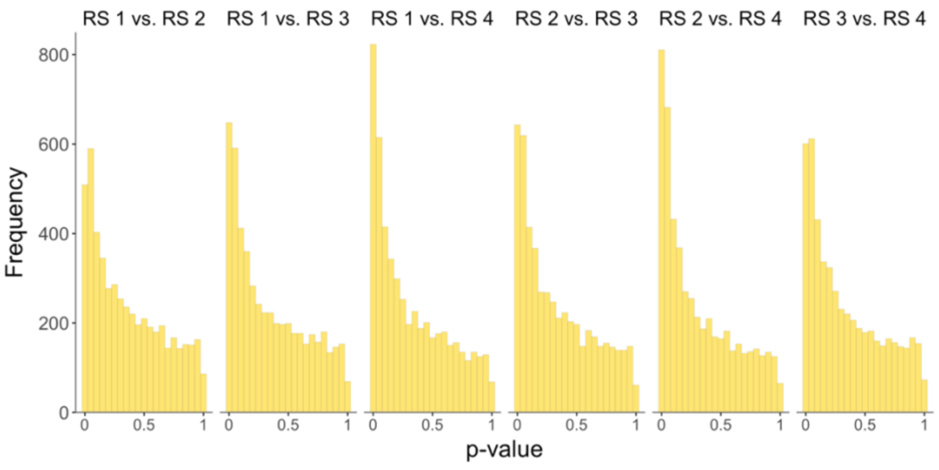


**Fig. S5.** p-value distributions of p-values obtained from differential methylation analysis. Pairwise comparison of the four reproductive stages are displayed in individual plots.

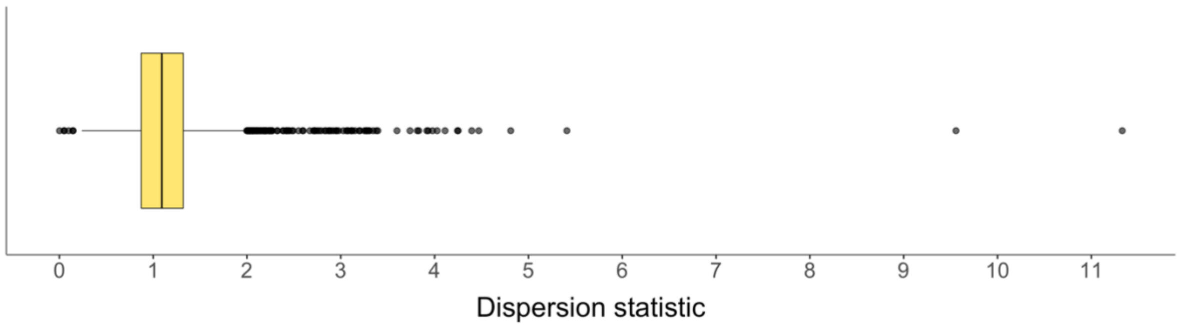


**Fig. S6.** Boxplot of dispersion statistics. Plot shows all CpG sites tested (including outliers) in the differential methylation analysis.

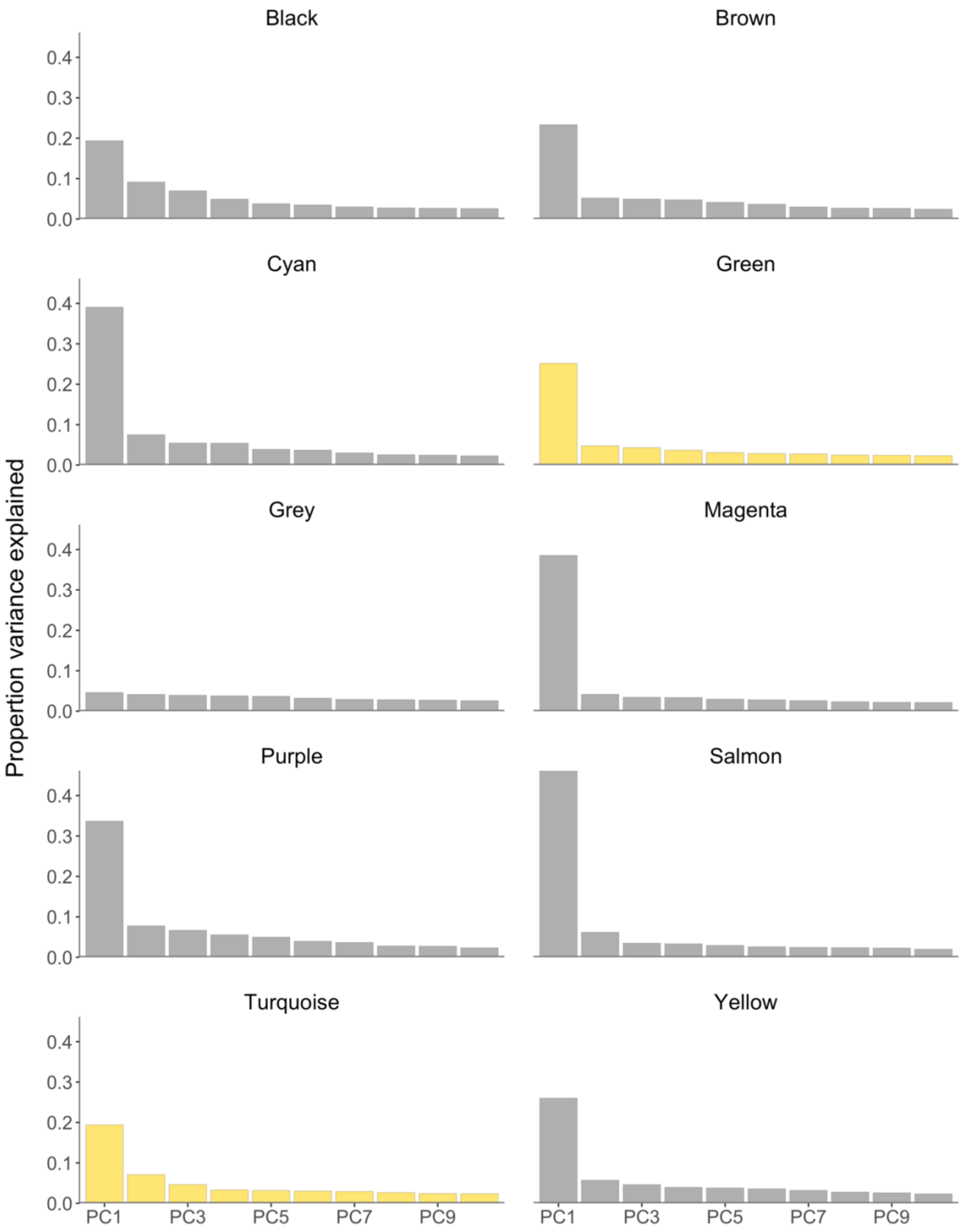


**Fig. S7.** Proportion of variance explained by the first ten principal components (PC1-PC10) for all modules found with weighted co-methylation network analysis. Modules (turquoise and green) with significant correlation to lay date centered sampling date are highlighted in yellow.

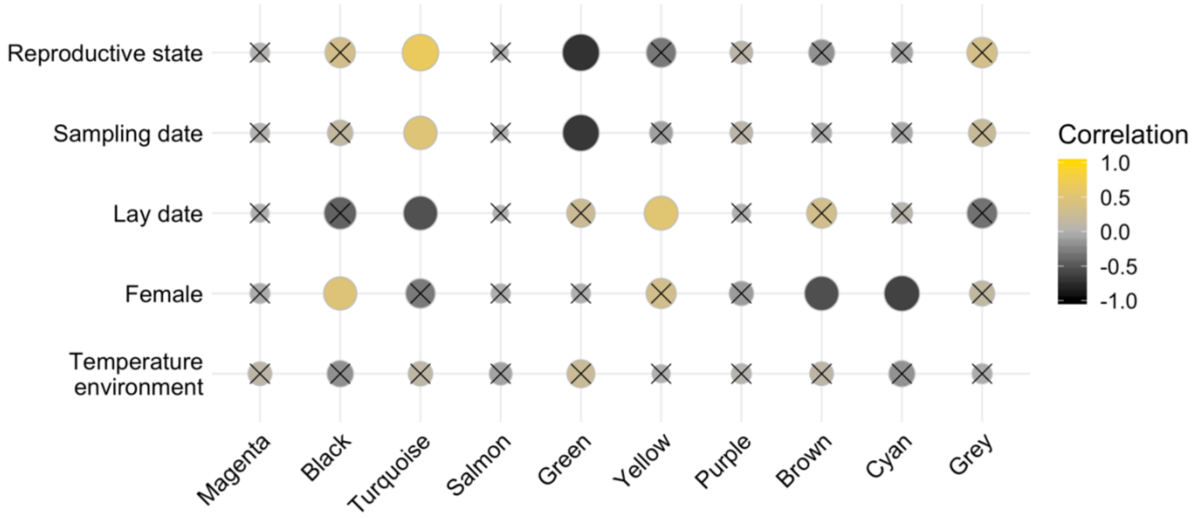


**Fig. S8.** Correlation of traits (reproductive state, sampling date, lay date, female, and temperature environment) with the module eigensite (i.e. first principal component). Color gradient indicates the direction of the correlation; positive correlation (yellow), no correlation (grey), and negative correlation (black). Size of spheres relates to the significance of the underlying correlation. Non-significant correlations (p-vale > 0.001, Bonferroni-corrected α-threshold) are crossed. See Supplementary Table 9 for correlation estimates and p-values.

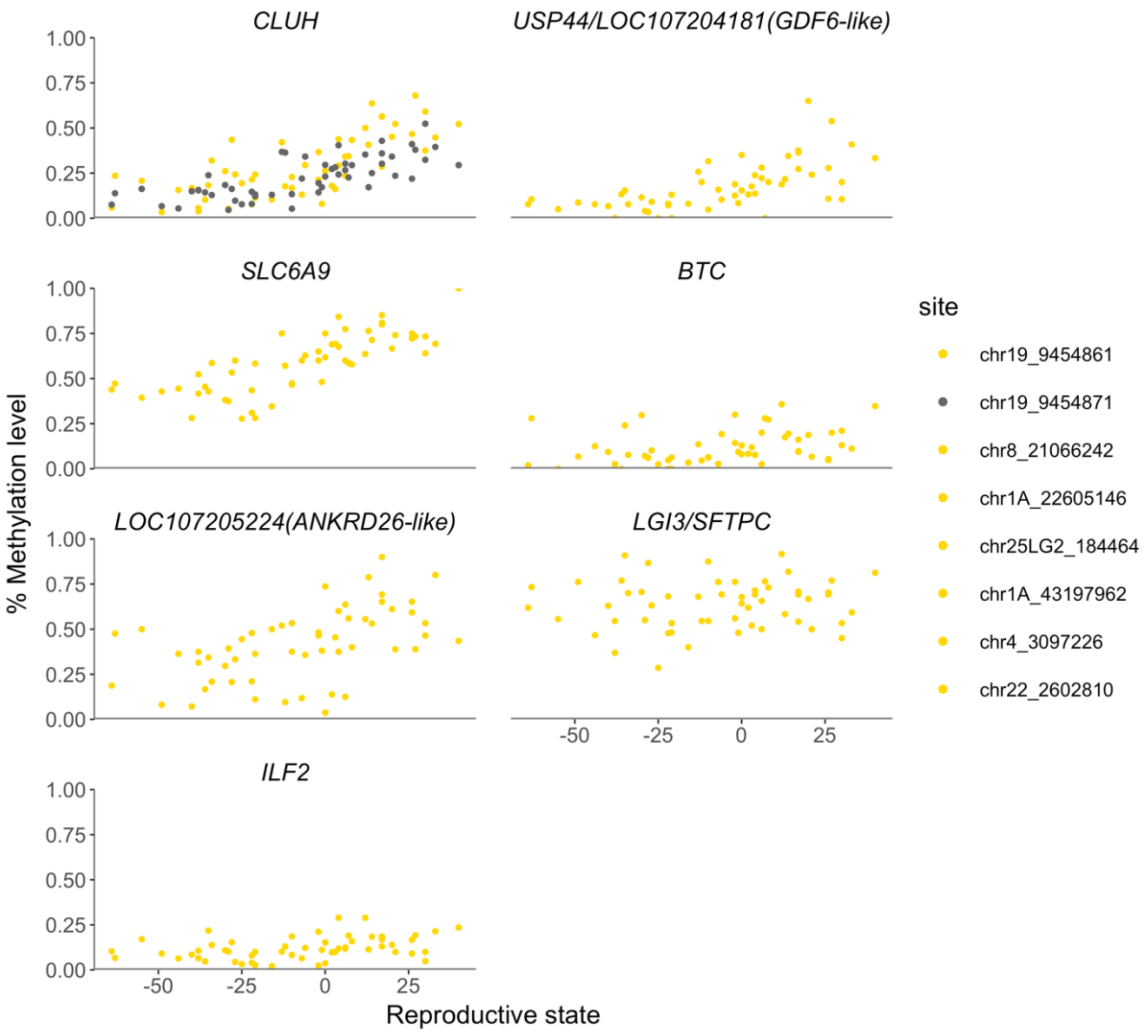


**Fig. S9.** Methylation profile of CpG sites that that showed significant variation in DNA methylation in the differential methylation analysis, but not in the co-methylation analysis. CpG sites were located within the regulatory region of *CLUH* (chr19), *SLC6A9* (chr8), *LOC107205224* (*ANKRD26*-like, chr1A), *ILF2* (chr25LG2), *USP44/LOC107204181* (*GDF6*-like, chr1A), *BTC* (chr4), and *LGI3/SFTPC* (chr22).

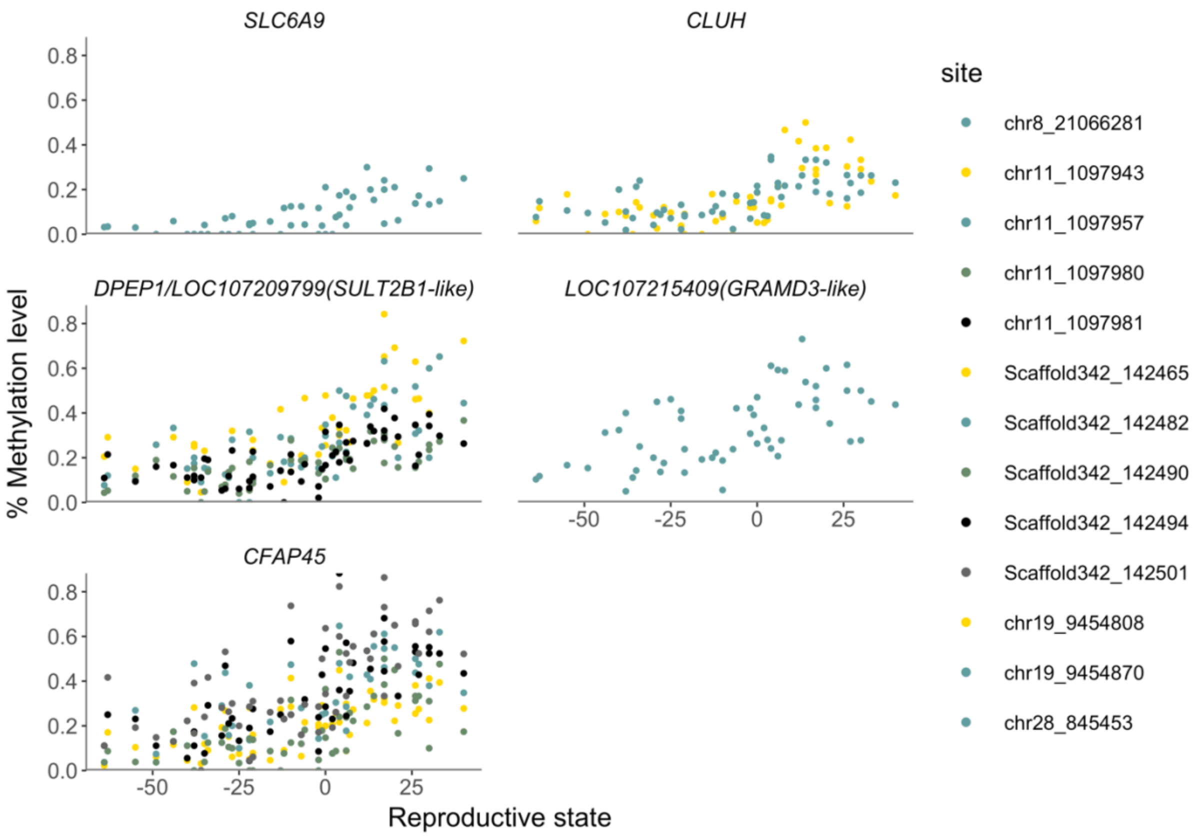


**Fig. S10.** Methylation profile of CpG sites that showed significant variation in DNA methylation in the co-methylation analysis (turquoise module), but not in the differential methylation analysis. CpG sites were located within the regulatory region of *SLC6A9* (chr8), *DPEP1/ LOC107209799 (SULT2B1*-like, chr11), *CFAP45* (Scaffold342), *CLUH* (chr19), LOC107215409 (*GRAMD3*-like, chr28).

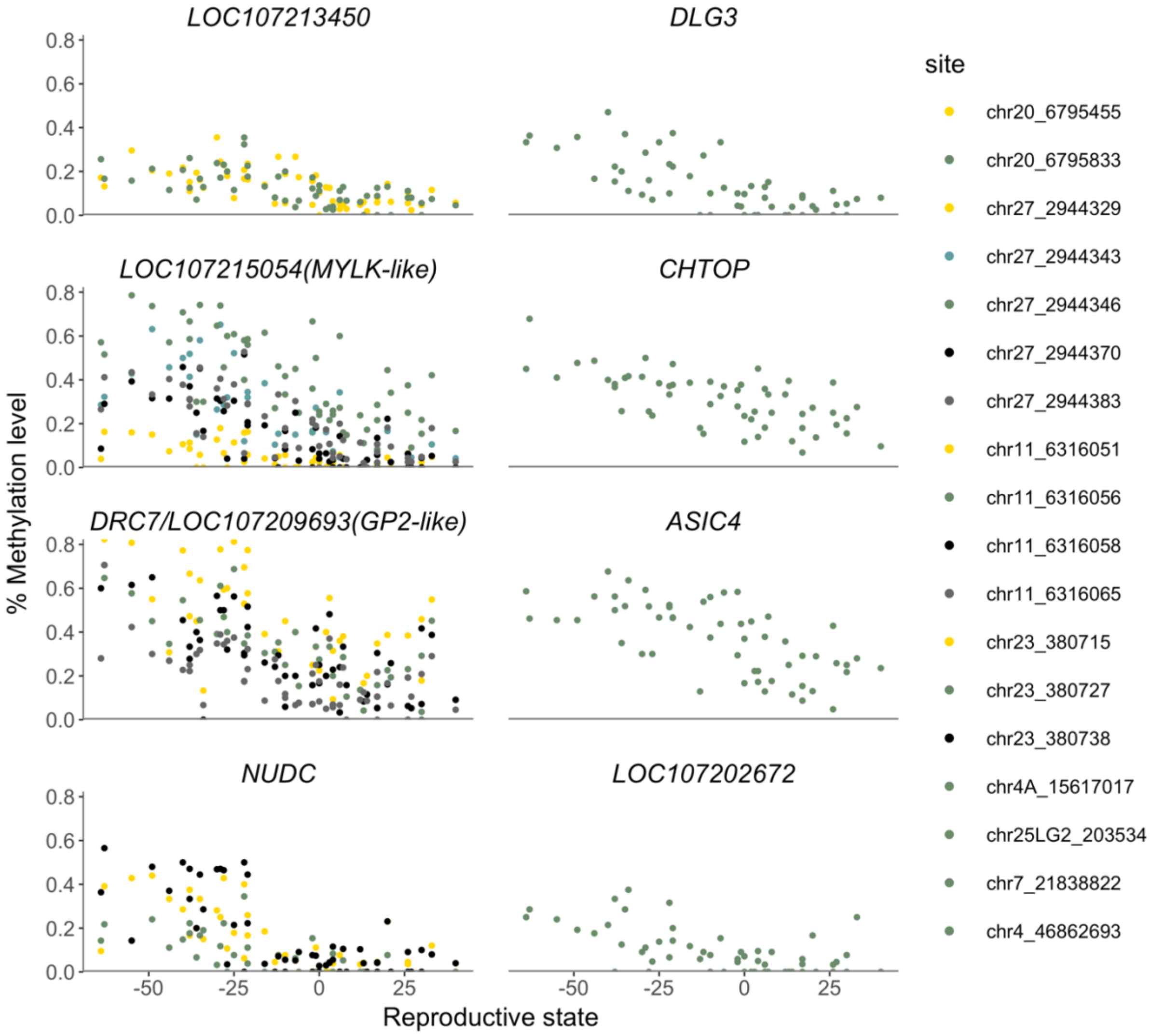


**Fig. S11.** Methylation profile of CpG sites that showed significant variation in DNA methylation in the co-methylation analysis (green module), but not in the differential methylation analysis. CpG sites were located within the regulatory region of *LOC107213450* (chr20), *LOC107215054* (*MYLK*-like, chr27), *DRC7*/ LOC107209693 (GP2-like, chr11), *NUDC* (chr23), *DLG3* (chr4A), *CHTOP* (chr25LG2), *ASIC4* (chr7), and *LOC107202672* (chr4).

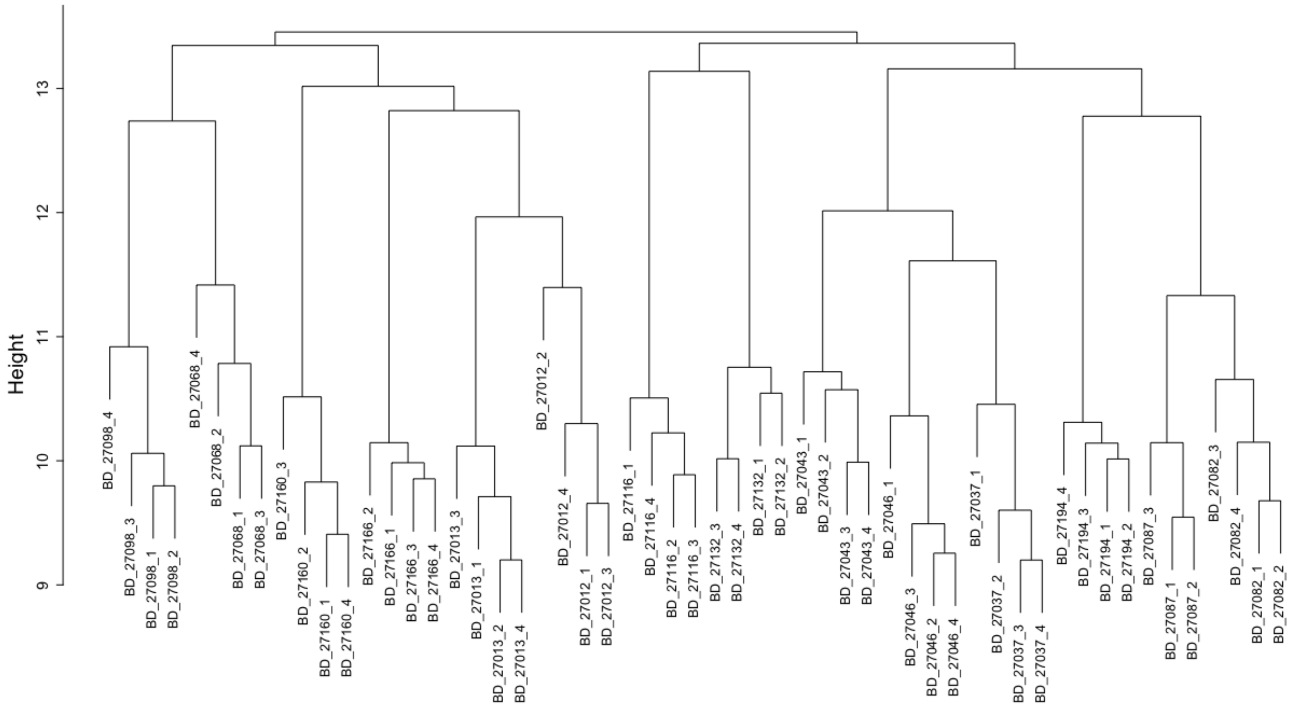


**Fig. S12.** Sample clustering to detect outliers.

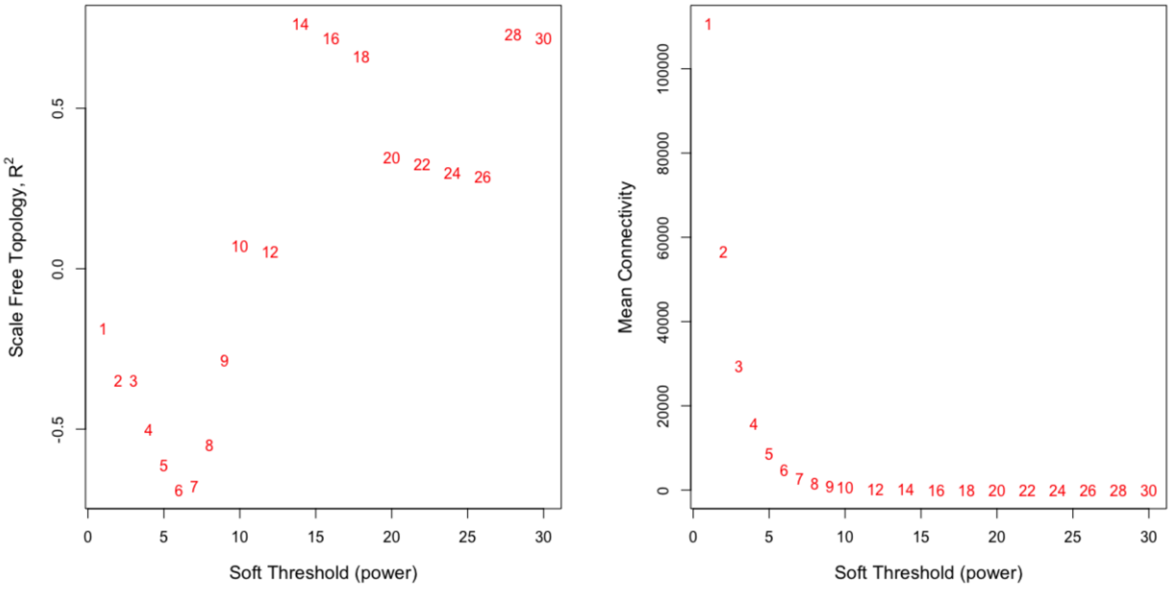


**Fig. S13.** Scale free topology (R2, left plot) and mean connectivity (right plot) for possible soft threshold values (power). Methylation profiles of all 223,282 CpG sites within the regulatory region of annotated genes were used.

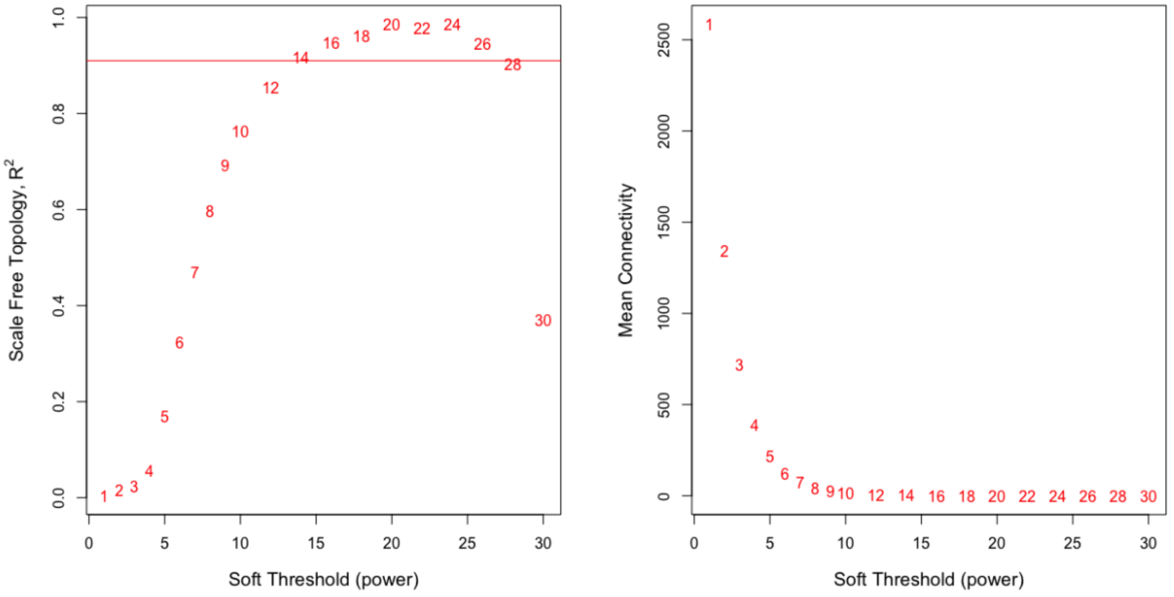


**Fig. S14.** Scale free topology (R2, left plot) and mean connectivity (right plot) for possible soft threshold values (power). Methylation profiles of the 5097 CpG sites also used in the differential methylation analysis were used.

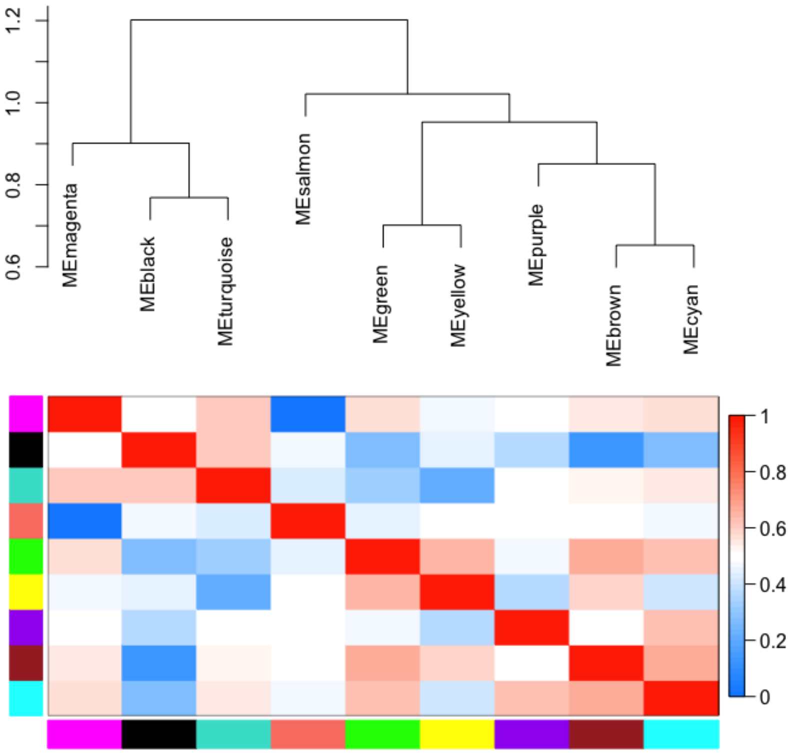


**Fig. S15.** Module eigensite clustering tree (top panel) and heatmap displaying adjacency between module eigensite (bottom panel). Please note that heatmap displays the adjacency between module eigensite which ranges from 0 (blue) to 1 (red). Here, merge-cut-height of 0.65 was used for module detection.

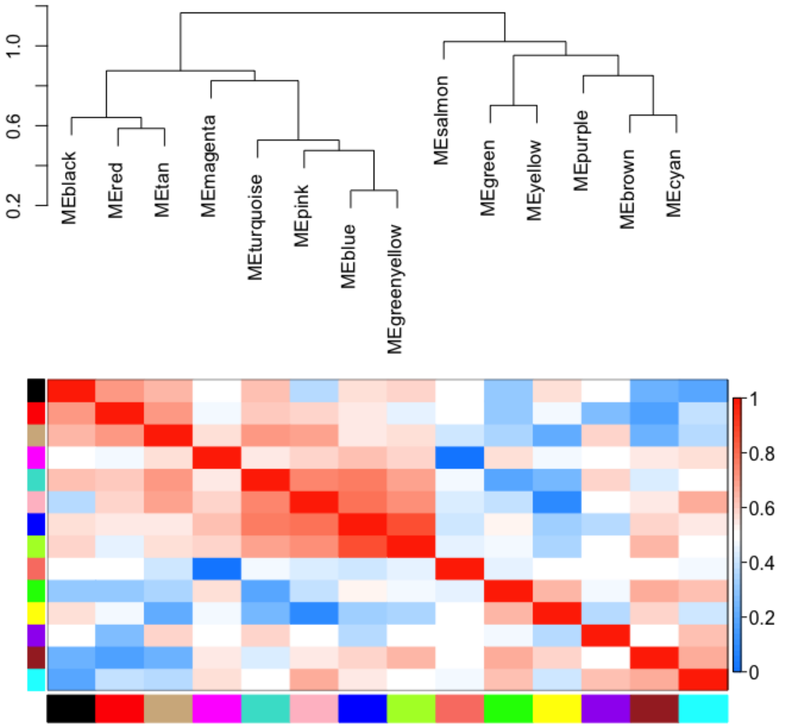


**Fig. S16.** Module eigensite clustering tree using (top panel) and heatmap displaying adjacency between module eigensite (bottom panel). Please note that heatmap displays the adjacency between module eigensite which ranges from 0 (blue) to 1 (red). Here, merge-cut-height of 0.15 was used for module detection.

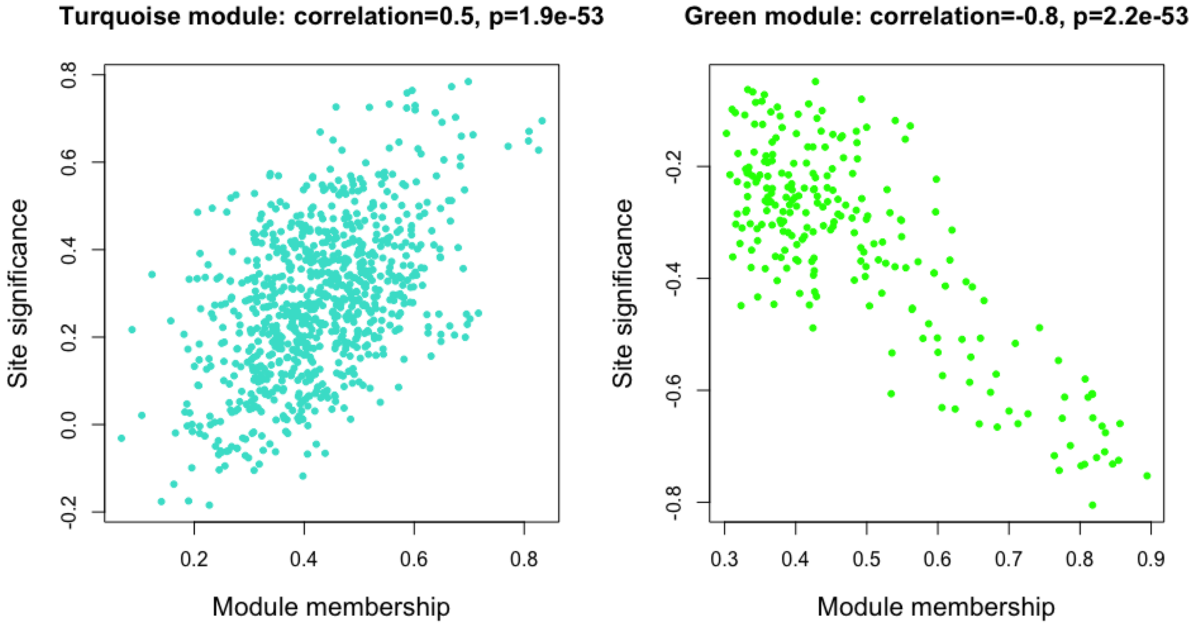


**Fig. S17.** Correlation between the trait-based site significance (SS) and module membership (MM) for the turquoise and green module. Both modules showed a significant correlation (of module 'eigensite') to the reproductive state See Supplementary Methods 5 for details on trait-based SS and MM.

SUPPLEMENTARY TABLES

**Table S1.** CpG sites that showed significant variation in DNA methylation between the second pre-laying stage and post-laying stage after correction for multiple testing (Bonferroni significance threshold: p-value < 9.81e-06) and filter for over- and under-dispersion of the model.

| Gene | Position | p-value | Dispersion statistic |
|---|---|---|---|
| *LOC107215054 (MYLK*-like*)* | chr27_2944354 | 6.97E-17 | 0.96 |
| *NR5A1* | chr17_1028562 | 3.32E-12 | 0.79 |
| *DRC7* | chr11_6316050 | 3.77E-10 | 0.82 |
| *LOC107209693 (GP2*-like*)* | chr11_6316050 | 3.77E-10 | 0.82 |
| *CFAP45* | Scaffold342_142514 | 1.45E-09 | 1.13 |
| *CLUH* | chr19_9454861 | 2.27E-09 | 0.91 |
| *NR5A1* | chr17_1028567 | 5.29E-09 | 1.00 |
| *LOC107215054 (MYLK*-like*)* | chr27_2944382 | 8.42E-09 | 1.25 |
| *DLG3* | chr4A_15617013 | 1.53E-08 | 1.12 |
| *LOC107213450* | chr20_6795879 | 1.98E-08 | 0.90 |
| *DRC7* | chr11_6316068 | 2.31E-08 | 0.80 |
| *LOC107209693 (GP2*-like*)* | chr11_6316068 | 2.31E-08 | 0.80 |
| *DRC7* | chr11_6316055 | 6.03E-08 | 0.95 |
| *LOC107209693 (GP2*-like*)* | chr11_6316055 | 6.03E-08 | 0.95 |
| *SLC6A9* | chr8_21066242 | 7.17E-08 | 1.26 |
| *DRC7* | chr11_6316057 | 2.65E-07 | 0.74 |
| *LOC107209693 (GP2*-like*)* | chr11_6316057 | 2.65E-07 | 0.74 |
| *CLUH* | chr19_9454871 | 4.37E-07 | 0.94 |
| *LOC107213450* | chr20_6795454 | 5.05E-07 | 1.02 |
| *SLC6A9* | chr8_21066275 | 6.19E-07 | 1.04 |
| *DRC7* | chr11_6316069 | 6.55E-07 | 0.75 |
| *LOC107209693 (GP2*-like*)* | chr11_6316069 | 6.55E-07 | 0.75 |
| *NR5A1* | chr17_1028571 | 1.04E-06 | 0.70 |
| *LOC107205224 (ANKRD26*-like*)* | chr1A_22605146 | 4.24E-06 | 1.15 |
| *ILF2* | chr25LG2_184464 | 6.38E-06 | 0.93 |
| *USP44* | chr1A_43197962 | 7.28E-06 | 1.08 |

| | | | |
|---|---|---|---|
| *LOC107204181 (GDF6*-like*)* | chr1A_43197962 | 7.28E-06 | 1.08 |
| *BTC* | chr4_3097226 | 8.23E-06 | 1.15 |
| *SFTPC* | chr22_2602810 | 9.81E-06 | 0.82 |
| *LGI3* | chr22_2602810 | 9.81E-06 | 0.82 |

**Table S2.** The most significant CpG site of each gene with at least one CpG site that showed significant variation in DNA methylation between the second pre-laying stage and post-laying stage ranked by the most significant CpG site.

| Gene Symbol | Position | p-value | Dispersion statistic | Rank |
|---|---|---|---|---|
| *LOC107215054 (MYLK*-like*)* | chr27_2944354 | 6.97E-17 | 0.96 | 1 |
| *NR5A1* | chr17_1028562 | 3.32E-12 | 0.79 | 2 |
| *DRC7* | chr11_6316050 | 3.77E-10 | 0.82 | 3 |
| *LOC107209693 (GP2*-like*)* | chr11_6316050 | 3.77E-10 | 0.82 | 3 |
| *CFAP45* | Scaffold342_142514 | 1.45E-09 | 1.13 | 4 |
| *CLUH* | chr19_9454861 | 2.27E-09 | 0.91 | 5 |
| *DLG3* | chr4A_15617013 | 1.53E-08 | 1.12 | 6 |
| *LOC107213450* | chr20_6795879 | 1.98E-08 | 0.90 | 7 |
| *SLC6A9* | chr8_21066242 | 7.17E-08 | 1.26 | 8 |
| *LOC107205224 (ANKRD26*-like*)* | chr1A_22605146 | 4.24E-06 | 1.15 | 9 |
| *ILF2* | chr25LG2_184464 | 6.38E-06 | 0.93 | 10 |
| *USP44* | chr1A_43197962 | 7.28E-06 | 1.08 | 11 |
| *LOC107204181 (GDF6*-like*)* | chr1A_43197962 | 7.28E-06 | 1.08 | 11 |
| *BTC* | chr4_3097226 | 8.23E-06 | 1.15 | 12 |
| *SFTPC* | chr22_2602810 | 9.81E-06 | 0.82 | 13 |
| *LGI3* | chr22_2602810 | 9.81E-06 | 0.82 | 13 |

**Table S3.** Number of CpG sites within each module. *CpG sites within the grey module did not cluster with other CpG sites.

| Module Color | Number of CpG sites |
|---|---|
| black | 426 |
| brown | 294 |
| cyan | 55 |
| green | 234 |
| grey* | 2750 |
| magenta | 105 |
| purple | 68 |

| | |
|---|---|
| salmon | 58 |
| turquoise | 826 |
| yellow | 281 |

**Table S4.** CpG sites with significant trait-based site significance (SS) and module membership (MM) within the turquoise module.

| Gene Symbol | Position | SS | p-value SS | MM | p-value MM |
|---|---|---|---|---|---|
| *NR5A1* | chr17_1028567 | 0.78 | 1.40E-12 | 0.70 | 3.08E-09 |
| *NR5A1* | chr17_1028562 | 0.77 | 4.77E-12 | 0.67 | 2.53E-08 |
| *SLC6A9* | chr8_21066275 | 0.73 | 2.65E-10 | 0.60 | 1.19E-06 |
| *NR5A1* | chr17_1028571 | 0.72 | 6.15E-10 | 0.60 | 1.18E-06 |
| *DPEP1* | chr11_1097980 | 0.71 | 1.01E-09 | 0.64 | 1.57E-07 |
| *LOC107209799 (SULT2B1*-like) | chr11_1097980 | 0.71 | 1.01E-09 | 0.64 | 1.57E-07 |
| *DPEP1* | chr11_1097943 | 0.70 | 2.21E-09 | 0.68 | 1.57E-08 |
| *LOC107209799 (SULT2B1*-like) | chr11_1097943 | 0.70 | 2.21E-09 | 0.68 | 1.57E-08 |
| *CFAP45* | Scaffold342_142494 | 0.69 | 4.04E-09 | 0.83 | 3.30E-15 |
| *SLC6A9* | chr8_21066281 | 0.69 | 5.16E-09 | 0.65 | 7.61E-08 |
| *CFAP45* | Scaffold342_142501 | 0.67 | 2.13E-08 | 0.81 | 8.14E-14 |
| *CFAP45* | Scaffold342_142514 | 0.66 | 3.57E-08 | 0.71 | 1.62E-09 |
| *DPEP1* | chr11_1097957 | 0.66 | 4.36E-08 | 0.69 | 7.64E-09 |
| *LOC107209799 (SULT2B1*-like) | chr11_1097957 | 0.66 | 4.36E-08 | 0.69 | 7.64E-09 |
| *CFAP45* | Scaffold342_142465 | 0.65 | 8.55E-08 | 0.81 | 9.06E-14 |
| *CFAP45* | Scaffold342_142482 | 0.64 | 1.79E-07 | 0.77 | 5.88E-12 |
| *DPEP1* | chr11_1097981 | 0.63 | 2.46E-07 | 0.61 | 9.02E-07 |
| *LOC107209799 (SULT2B1*-like) | chr11_1097981 | 0.63 | 2.46E-07 | 0.61 | 9.02E-07 |
| *CFAP45* | Scaffold342_142490 | 0.63 | 2.96E-07 | 0.83 | 8.15E-15 |
| *CLUH* | chr19_9454808 | 0.62 | 4.69E-07 | 0.61 | 6.74E-07 |
| *LOC107215409 (GRAMD3*-like) | chr28_845453 | 0.61 | 7.14E-07 | 0.68 | 8.39E-09 |
| *CLUH* | chr19_9454870 | 0.61 | 9.82E-07 | 0.65 | 6.16E-08 |

**Table S5.** CpG sites with significant trait-based site significance (SS) and module membership (MM) within the green module.

| Gene Symbol | Position | SS | p-value SS | MM | p-value MM |
|---|---|---|---|---|---|
| *LOC107213450* | chr20_6795879 | -0.81 | 1.28E-13 | 0.82 | 2.58E-14 |
| *LOC107215054* (*MYLK*-like) | chr27_2944383 | -0.75 | 3.41E-11 | 0.89 | 3.68E-20 |
| *DRC7* | chr11_6316068 | -0.74 | 8.23E-11 | 0.77 | 5.89E-12 |
| *LOC107209693* (*GP2*-like) | chr11_6316068 | -0.74 | 8.23E-11 | 0.77 | 5.89E-12 |
| *NUDC* | chr23_380715 | -0.73 | 1.69E-10 | 0.80 | 1.99E-13 |
| *LOC107213450* | chr20_6795454 | -0.73 | 2.13E-10 | 0.81 | 1.09E-13 |
| *LOC107215054* (*MYLK*-like) | chr27_2944343 | -0.73 | 2.23E-10 | 0.85 | 4.37E-16 |
| *DRC7* | chr11_6316050 | -0.73 | 3.85E-10 | 0.85 | 1.10E-16 |
| *LOC107209693* (*GP2*-like) | chr11_6316050 | -0.73 | 3.85E-10 | 0.85 | 1.10E-16 |
| *LOC107215054* (*MYLK*-like) | chr27_2944346 | -0.72 | 5.68E-10 | 0.82 | 1.21E-14 |
| *NUDC* | chr23_380738 | -0.72 | 7.51E-10 | 0.76 | 1.17E-11 |
| *DLG3* | chr4A_15617017 | -0.71 | 1.29E-09 | 0.83 | 2.43E-15 |
| *LOC107215054* (*MYLK*-like) | chr27_2944370 | -0.70 | 2.98E-09 | 0.79 | 1.13E-12 |
| *DLG3* | chr4A_15617013 | -0.68 | 1.52E-08 | 0.84 | 2.06E-15 |
| *CHTOP* | chr25LG2_203534 | -0.67 | 2.94E-08 | 0.68 | 8.85E-09 |
| *LOC107215054* (*MYLK*-like) | chr27_2944354 | -0.66 | 3.27E-08 | 0.83 | 4.14E-15 |
| *ASIC4* | chr7_21838822 | -0.66 | 4.22E-08 | 0.66 | 4.74E-08 |
| *NUDC* | chr23_380727 | -0.66 | 4.30E-08 | 0.71 | 1.03E-09 |
| *DRC7* | chr11_6316051 | -0.66 | 4.36E-08 | 0.86 | 7.97E-17 |
| *LOC107209693* (*GP2*-like) | chr11_6316051 | -0.66 | 4.36E-08 | 0.86 | 7.97E-17 |
| *DRC7* | chr11_6316055 | -0.65 | 7.88E-08 | 0.77 | 3.82E-12 |
| *LOC107209693* (*GP2*-like) | chr11_6316055 | -0.65 | 7.88E-08 | 0.77 | 3.82E-12 |
| *DRC7* | chr11_6316069 | -0.65 | 8.31E-08 | 0.82 | 2.47E-14 |
| *LOC107209693* (*GP2*-like) | chr11_6316069 | -0.65 | 8.31E-08 | 0.82 | 2.47E-14 |
| *DRC7* | chr11_6316058 | -0.64 | 1.27E-07 | 0.73 | 3.33E-10 |
| *LOC107209693* (*GP2*-like) | chr11_6316058 | -0.64 | 1.27E-07 | 0.73 | 3.33E-10 |
| *DRC7* | chr11_6316065 | -0.64 | 1.70E-07 | 0.70 | 2.68E-09 |
| *LOC107209693* (*GP2*-like) | chr11_6316065 | -0.64 | 1.70E-07 | 0.70 | 2.68E-09 |

| | | | | | |
|---|---|---|---|---|---|
| *LOC107213450* | chr20_6795455 | -0.63 | 2.10E-07 | 0.62 | 3.51E-07 |
| *LOC107213450* | chr20_6795833 | -0.63 | 2.44E-07 | 0.61 | 9.55E-07 |
| *DRC7* | chr11_6316056 | -0.61 | 6.71E-07 | 0.81 | 5.94E-14 |
| *LOC107209693 (GP2*-like*)* | chr11_6316056 | -0.61 | 6.71E-07 | 0.81 | 5.94E-14 |
| *DRC7* | chr11_6316057 | -0.61 | 6.85E-07 | 0.78 | 2.66E-12 |
| *LOC107209693 (GP2*-like*)* | chr11_6316057 | -0.61 | 6.85E-07 | 0.78 | 2.66E-12 |
| *LOC107215054 (MYLK*-like*)* | chr27_2944382 | -0.61 | 8.89E-07 | 0.82 | 2.61E-14 |
| *LOC107215054 (MYLK*-like*)* | chr27_2944329 | -0.61 | 9.46E-07 | 0.82 | 2.81E-14 |
| *LOC107202672* | chr4_46862693 | -0.60 | 1.07E-06 | 0.67 | 1.69E-08 |

**Table S6.** The most significant CpG site of each gene with at least one CpG sites with significant trait-based site significance (SS) and module membership (MM) within the turquoise module ranked by the most significant CpG site.

| Gene Symbol | Position | SS | p-value SS | MM | p-value MM | Rank |
|---|---|---|---|---|---|---|
| *NR5A1* | chr17_1028567 | 0.78 | 1.40E-12 | 0.70 | 3.08E-09 | 1 |
| *SLC6A9* | chr8_21066275 | 0.73 | 2.65E-10 | 0.60 | 1.19E-06 | 2 |
| *DPEP1* | chr11_1097980 | 0.71 | 1.01E-09 | 0.64 | 1.57E-07 | 3 |
| *LOC107209799 (SULT2B1*-like*)* | chr11_1097980 | 0.71 | 1.01E-09 | 0.64 | 1.57E-07 | 3 |
| *CFAP45* | Scaffold342_142494 | 0.69 | 4.04E-09 | 0.83 | 3.30E-15 | 4 |
| *CLUH* | chr19_9454808 | 0.62 | 4.69E-07 | 0.61 | 6.74E-07 | 5 |
| *LOC107215409 (GRAMD3*-like*)* | chr28_845453 | 0.61 | 7.14E-07 | 0.68 | 8.39E-09 | 6 |

**Table S7.** The most significant CpG site of each gene with at least one CpG sites with significant trait-based site significance (SS) and module membership (MM) within the green module ranked by the most significant CpG site.

| Gene Symbol | Position | SS | p-value SS | MM | p-value MM | Rank |
|---|---|---|---|---|---|---|
| *LOC107213450* | chr20_6795879 | -0.81 | 1.28E-13 | 0.82 | 2.58E-14 | 1 |
| *LOC107215054 (MYLK*-like*)* | chr27_2944383 | -0.75 | 3.41E-11 | 0.89 | 3.68E-20 | 2 |
| *DRC7* | chr11_6316068 | -0.74 | 8.23E-11 | 0.77 | 5.89E-12 | 3 |
| *LOC107209693 (GP2*-like*)* | chr11_6316068 | -0.74 | 8.23E-11 | 0.77 | 5.89E-12 | 3 |
| *NUDC* | chr23_380715 | -0.73 | 1.69E-10 | 0.80 | 1.99E-13 | 4 |
| *DLG3* | chr4A_15617017 | -0.71 | 1.29E-09 | 0.83 | 2.43E-15 | 5 |
| *CHTOP* | chr25LG2_203534 | -0.67 | 2.94E-08 | 0.68 | 8.85E-09 | 6 |
| *ASIC4* | chr7_21838822 | -0.66 | 4.22E-08 | 0.66 | 4.74E-08 | 7 |
| *LOC107202672* | chr4_46862693 | -0.60 | 1.07E-06 | 0.67 | 1.69E-08 | 8 |

**Table S8.** Significant CpG sites found with the differential methylation analysis (DMA) and the co-methylation analysis (WGCNA) ranked by their sum of ranks (i.e. rank DMA + rank WGCNA).

| Gene Symbol | Position | Module Color | Rank DMA | Rank WGCNA | Sum of Ranks |
|---|---|---|---|---|---|
| *NR5A1* | chr17_1028562 | turquoise | 2 | 1 | 3 |
| *NR5A1* | chr17_1028567 | turquoise | 2 | 1 | 3 |
| *NR5A1* | chr17_1028571 | turquoise | 2 | 1 | 3 |
| *LOC107215054 (MYLK*-like*)* | chr27_2944354 | green | 1 | 2 | 3 |
| *LOC107215054 (MYLK*-like*)* | chr27_2944382 | green | 1 | 2 | 3 |
| *DRC7* | chr11_6316050 | green | 3 | 3 | 6 |
| *LOC107209693 (GP2*-like*)* | chr11_6316050 | green | 3 | 3 | 6 |
| *DRC7* | chr11_6316055 | green | 3 | 3 | 6 |
| *LOC107209693 (GP2*-like*)* | chr11_6316055 | green | 3 | 3 | 6 |
| *DRC7* | chr11_6316057 | green | 3 | 3 | 6 |
| *LOC107209693 (GP2*-like*)* | chr11_6316057 | green | 3 | 3 | 6 |
| *DRC7* | chr11_6316068 | green | 3 | 3 | 6 |
| *LOC107209693 (GP2*-like*)* | chr11_6316068 | green | 3 | 3 | 6 |
| *DRC7* | chr11_6316069 | green | 3 | 3 | 6 |
| *LOC107209693 (GP2*-like*)* | chr11_6316069 | green | 3 | 3 | 6 |
| *LOC107213450* | chr20_6795454 | green | 7 | 1 | 8 |
| *LOC107213450* | chr20_6795879 | green | 7 | 1 | 8 |
| *CFAP45* | Scaffold342_142514 | turquoise | 4 | 4 | 8 |
| *SLC6A9* | chr8_21066275 | turquoise | 8 | 2 | 10 |
| *DLG3* | chr4A_15617013 | green | 6 | 5 | 11 |

**Table S9.** Significant correlations between module eigensites and traits (reproductive state, sampling date, laying date, and female Id).

| Module | Trait | Correlation | p-value |
|---|---|---|---|
| turquoise | Reproductive state | 0.66 | 3.15E-08 |
| green | Reproductive state | -0.71 | 9.63E-10 |
| turquoise | Sampling date | 0.48 | 2.16E-04 |
| green | Sampling date | -0.68 | 1.32E-08 |

| | | | |
|---|---|---|---|
| turquoise | Laying date | -0.53 | 2.71E-05 |
| yellow | Laying date | 0.51 | 7.32E-05 |
| black | Female Id | 0.47 | 2.48E-04 |
| brown | Female Id | -0.54 | 2.53E-05 |
| cyan | Female Id | -0.61 | 8.16E-07 |

**Table S10.** Gene lists for GO analysis. Gene lists include genes with CpG sites that show significant variation in DNA methylation in the differential methylation analysis (DMA) or co-methylation analysis (WGCNA). The background list includes all genes that have at least one CpG site within their regulatory region.

| Analysis | Gene List | Number of Genes |
|---|---|---|
| DMA | Second pre-laying vs. post-laying stage | 16 |
| DMA | Any pairwise comparison | 35 |
| WGCNA | Turquoise module | 7 |
| WGCNA | Green module | 9 |
| WGCNA | Turquoise and green module | 16 |
| -NA- | Background | 12324 |
